# Heat-Induced Phosphatidylserine Changes Drive HSPA1A’s Plasma Membrane Localization

**DOI:** 10.1101/2024.12.02.626454

**Authors:** Jensen Low, Rachel Altman, Allen Badolian, Azalea Blythe Cuaresma, Carolina Briseño, Uri Keshet, Oliver Fiehn, Robert V. Stahelin, Nikolas Nikolaidis

## Abstract

Heat shock protein A1A (HSPA1A) is a molecular chaperone crucial in cell survival. In addition to its cytosolic functions, HSPA1A translocates to heat-shocked and cancer cells’ plasma membrane (PM). In cancer, PM-localized HSPA1A (mHSPA1A) is associated with increased tumor aggressiveness and therapeutic resistance, suggesting that preventing its membrane localization could have therapeutic value. This translocation depends on HSPA1A’s interaction with PM phospholipids, including phosphatidylserine (PS). Although PS binding regulates HSPA1A’s membrane localization, the exact trigger for this movement remains unclear. Given that lipid modifications are a cancer hallmark, we hypothesized that PS is a crucial lipid driving HSPA1A translocation and that heat-induced changes in PS levels trigger HSPA1A’s PM localization in response to heat stress. We tested this hypothesis using pharmacological inhibition and RNA interference (RNAi) targeting PS synthesis, combined with confocal microscopy, lipidomics, and western blotting. Lipidomic analysis and PS-specific biosensors confirmed a heat shock-induced PS increase, peaking immediately post-stress. Inhibition of PS synthesis with fendiline and RNAi significantly reduced HSPA1A’s PM localization, while depletion of cholesterol or fatty acids had minimal effects, confirming specificity for PS. Further experiments showed that PS saturation and elongation changes did not significantly impact HSPA1A’s PM localization, indicating that the total PS increase, rather than specific PS species, is the critical factor. These findings reshape current models of HSPA1A trafficking, demonstrating that PS is a crucial regulator of HSPA1A’s membrane translocation during the heat shock response. This work offers new insights into lipid-regulated protein trafficking and highlights the importance of PS in controlling cellular responses to stress.

## Introduction

Heat shock protein A1A (HSPA1A) is a 70-kDa molecular chaperone essential for the cellular stress response(1,2). Primarily functioning in the cytosol, HSPA1A also translocates to the plasma membrane (PM) of heat-shocked and cancer cells, where its PM-localized form (mHSPA1A) is associated with increased tumor aggressiveness and therapeutic resistance (3–10).

Several studies revealed that although HSPA1A lacks canonical lipid-binding domains, it associates with cellular membranes and liposomes rich in anionic lipids, particularly phosphatidylserine (PS), while avoiding neutral lipid environments like those composed only of phosphatidylcholine (PC)(6,9–19). HSPA1A’s interaction with PS is driven by a combination of electrostatic and hydrophobic forces rather than the general anionic charge of the membrane (3,15–18,20–22). Notably, HSPA1A’s embedding in PS-enriched liposomes is promoted by high PS saturation—an alteration commonly observed in cancer and stressed cells (13,15,16,23–28). Given this, the finding that mHSPA1A localization is critically dependent on PS interaction, though significant, is not unexpected.

Recent research suggests that HSPA1A’s translocation to the plasma membrane in response to stress is triggered by binding to intracellular PS (6,9,11,14). Following heat shock and during recovery, HSPA1A integrates into the PM and can be released into the extracellular space in a membrane-bound form. This PS-mediated translocation enables key biological functions of membrane-associated HSPA1A, including immune modulation, clathrin-independent endocytosis, viral entry facilitation, tumor cell survival, membrane stabilization, membrane chaperone activity, microautophagy, and signal transduction (3,29–31). Thus, HSPA1A’s interaction with PS may be essential in cancer progression, cellular stress responses, and other disease states.

Although HspA1A’s selective interaction with PS is essential for eliciting its movement from the cytosol to the plasma membrane in response to heat stress, the molecular triggers eliciting this association remain undefined. Cancer and heat-shocked cells commonly exhibit altered lipid composition (23–25), thus, we hypothesized that heat stress-driven increases in PS levels are a critical trigger for HSPA1A’s PM localization. Specifically, we proposed that heat stress modifies membrane lipids to enhance HSPA1A’s selective binding to PS-rich microdomains, thereby facilitating translocation from the cytosol to the plasma membrane.

To test this hypothesis, we characterized cellular lipid content after heat shock and during recovery. We examined the relationship between PS levels and HSPA1A’s translocation to the PM using pharmacological inhibition, RNA interference (RNAi), confocal microscopy, lipidomics, and western blotting.

## Materials and Methods

### Plasmids

To study HSPA1A localization, we utilized differentially tagged protein versions previously described in our earlier publications. Specifically, the mouse *hspa1a* cDNA sequence (accession number BC054782) was used to generate the recombinant constructs employed in this study. HSPA1A was tagged with green fluorescent protein (GFP), red fluorescent protein (RFP), or Myc epitope. The gene was subcloned into the pEGFP-C2 (producing N-terminally tagged HSPA1A), mRFP-C1 (producing N-terminally tagged HSPA1A), or pcDNA™3.1/myc-His(-) (producing C-terminally tagged HSPA1A) vectors for expression in mammalian cells. Subcloning was performed via directional cloning after PCR amplification and restriction enzyme digestion, following protocols detailed in the references (18) and (32). The phosphatidylserine (PS) biosensor Lact-C2-GFP and Lact-C2-mCherry plasmids were generously provided by Sergio Grinstein (Addgene plasmids #22852 and #17274, respectively) (33).

### Cell Culture

To examine HSPA1A localization, we utilized two human cell lines: human embryonic kidney cells (HEK293; ATCC® CRL-1573™) and HeLa cells derived from Henrietta Lacks (ATCC® CCL-2™), both obtained from ATCC in December 2016 and verified biannually. HEK293 cells were cultured in Dulbecco’s Modified Eagle Medium (DMEM), while HeLa cells were grown in Minimum Essential Medium (MEM). Media for both cell lines were supplemented with 10% fetal bovine serum (FBS), 2 mM L-glutamine, and penicillin-streptomycin. HeLa medium was supplemented with 0.1 mM non-essential amino acids (NEAA) and 1 mM sodium pyruvate. All cultures were maintained in a humidified atmosphere containing 5% CO₂ at 37°C.

### Transient Cell Transfections

#### Plasmid Transfections

Transient transfections were performed to express the proteins used in this study. Cells were seeded one day before transfection into either 24-well plates (2.0 × 10⁴ cells per well, with poly-D-lysine-treated coverslips) or 75 cm² cell culture flasks (6.0 × 10⁶ cells per flask). After 18 hours, cells were transfected with the appropriate construct using the PolyJet In Vitro DNA Transfection Reagent (SignaGen Laboratories, Frederick, MD, USA), following the manufacturer’s instructions. Transfection proceeded for 18 hours, after which the media was replaced with fresh, complete media to support subsequent experiments.

#### RNAi Transfections

To investigate the role of phosphatidylserine (PS) synthesis enzymes in HSPA1A’s plasma membrane (PM) localization, RNA interference (RNAi) was used to inhibit phosphatidylserine synthase enzymes 1 and 2 (PSS1, PSS2). The impact of these genetic interventions on HSPA1A localization and protein levels was assessed using confocal microscopy. Non-targeting control siRNA (Stealth^TM^ RNAi Negative control duplexes, Cat. No. 12935-300; Invitrogen, Carlsbad, CA, USA) was used as a negative control to account for off-target effects of RNAi.

For all RNAi transfections, cells were co-transfected with 0.75 µg of DNA encoding HSPA1A-GFP or Lact-C2-GFP. Experimental cells were transfected with 1.50 pmol of PSS1 RNAi (PTDSS1 Stealth siRNA, Cat. No. 1299001, assay ID: HSS114755; Invitrogen, Carlsbad, CA, USA), PSS2 RNAi (PTDSS2 Stealth siRNA Cat. No. 1299001, assay ID: HSS129844; Invitrogen, Carlsbad, CA, USA), or a combination of both, while control cells received 0.25 nM of non-targeting control siRNA. Transfections were carried out using the Lipofectamine™ 3000 Transfection Reagent, prepared by incubating the RNAi/DNA mixture with the reagent for 15 minutes at room temperature. This mixture was added to cells seeded on poly-D-lysine-coated coverslips in 24-well plates. Transfection proceeded for 24 hours, followed by the replacement of transfection media with fresh, complete media.

### Cell Treatments

#### Heat Shock Treatment

To assess the effect of heat shock on HSPA1A relocalization, cells were either maintained at 37°C or subjected to heat stress by incubating in a humidified CO₂ incubator equilibrated at 42°C for 60 minutes. Following heat shock, cells were allowed to recover at 37°C for 8 hours.

#### PS Inhibition

To investigate the impact of PS reduction on HSPA1A PM localization, cells were treated with fendiline (Cayman Chemical, Ann Arbor, MI, USA), an FDA-approved drug known to decrease PS content in MDCK (34) and HEK293 cells (35). Cells were exposed to either DMSO (control) or 10 µM fendiline diluted in serum-free MEM for 48 hours at 37°C in a humidified CO₂ incubator. Fendiline was added directly to the existing culture media without prior aspiration. Heat shock and recovery protocols were synchronized to conclude at the end of the 48-hour treatment.

#### Lipid Desaturation Inhibition

To examine the role of lipid desaturation in HSPA1A PM localization, cells were treated with desaturase inhibitors CP-24879 (HCl; Cayman Chemical, Ann Arbor, MI, USA) and SC 26196 (Santa Cruz Biotechnology, Dallas, TX, USA). CP-24879 is a mixed Δ5/Δ6 desaturase inhibitor tested in cell culture and animals (36,37), while SC 26196 selectively inhibits Δ6 desaturase with minimal effects on Δ5 and Δ9 desaturases (38,39). Cells were treated with either DMSO (control), 5.5 µM CP-24879, or 100 µM SC 26196 diluted in serum-free MEM for 48 hours at 37°C in a humidified CO₂ incubator. Drugs were added directly to the culture media, and heat shock and recovery were timed to conclude with the 48-hour treatment period.

#### Fatty Acid Synthesis Inhibition

To evaluate the effect of fatty acid synthesis on HSPA1A PM localization, cells were treated with cerulenin (Sigma-Aldrich, St. Louis, MO, USA), a fatty acid synthase (FASN) inhibitor (40). Cells were incubated with either DMSO (control) or 10 µg/ml cerulenin diluted in serum-free MEM for 9 hours at 37°C in a humidified CO₂ incubator. Cerulenin was added directly to the media without aspiration, and heat shock and recovery were completed by the end of the 9-hour treatment.

#### Cholesterol Depletion

To determine the relationship between HSPA1A PM localization and PM cholesterol, cells were treated with methyl-β-cyclodextrin (MβCD; Sigma-Aldrich, St. Louis, MO, USA), which depletes PM cholesterol and alters cholesterol domains on the cell surface (41). Cells were incubated with either DMSO (control) or 5 mM MβCD diluted in serum-free MEM for 9 hours at 37°C in a humidified CO₂ incubator. MβCD was directly added to the media, and heat shock and recovery protocols concluded at the end of the 9-hour treatment period.

#### Cell Viability Assay

Cell viability following heat shock and other treatments was assessed using the trypan blue exclusion assay. Viable and non-viable cells were quantified with the Cellometer® Auto X4 Cell Counter (Nexcelom Bioscience, Lawrence, MA, USA).

### Confocal Microscopy and Image Analysis

#### Cell Fixation and Staining

After transfection and subsequent treatments, cells were fixed in 4% paraformaldehyde (PFA) prepared in a complete growth medium for 12 minutes at room temperature (RT). Following fixation, PFA was aspirated, and the cells were washed three times with 1X phosphate-buffered saline (PBS). To stain the plasma membrane (PM), cells were incubated with 1 μg/ml wheat germ agglutinin (WGA) Alexa Fluor® 555 conjugate (Invitrogen, Carlsbad, CA, USA for 10 minutes at RT. After staining, the cells were washed three additional times with 1X PBS. Coverslips were mounted onto slides using approximately 3 μL of DAPI Fluoromount-G® (SouthernBiotech, Birmingham, AL, USA) and allowed to dry for 24 hours at RT in the dark. Prepared slides were stored at 4°C until visualization by confocal microscopy (19).

#### Microscope Setup

Cell imaging was performed using an Olympus FLUOVIEW FV3000 inverted scanning confocal microscope with a 60x 1.5 NA oil immersion objective. Multichannel image acquisition was conducted with the following settings: Channel 1 (Green): Excitation at 488 nm; emission at 510 nm; Channel 2 (Blue): Excitation at 405 nm; emission at 461 nm; Channel 3 (Red): Excitation at 561 nm; emission at 583 nm.

#### Image Analysis

Quantitative analysis was conducted on images from several cells (details provided in figure legends) collected across three independent experiments. Images were processed manually using ImageJ software (42), employing the corrected total cell fluorescence (CTCF) method (43). The formula used for CTCF was: CTCF = Integrated Density – (Area of Region of Interest * Fluorescence of background reading).

CTCF values were determined for both the PM and the cytosol. A ratio of PM fluorescence to total cell fluorescence (PM + cytosol) was calculated for each cell, following established protocols (32,44–46). As cell slices are not entirely flat, and the CTCF method utilizes the outermost pixels of each cell, a PM localization value of 0% cannot be achieved (44,45).

### Cell Surface Biotinylation

To further investigate the plasma membrane (PM) localization of HSPA1A, we performed cell surface biotinylation as described previously (19,45).

#### Biotinylation Procedure

Immediately after heat shock treatment, HEK293 cells were trypsinized, washed three times with PBS (pH 8.0), counted, and incubated in freshly prepared PBS (pH 8.0) containing 10 mM CaCl₂ and 1 mg/ml Sulfo-NHS-LC-biotin (Thermo Scientific Pierce, Rockford, IL, USA) for 30 minutes at room temperature with constant agitation. The biotinylation reaction was quenched by adding 100 mM glycine in PBS (pH 8.0) for 10 minutes.

#### Lysis and Streptavidin Pull-Down

Cells were lysed in 500 μl of radioimmunoprecipitation assay (RIPA) buffer. A total of 500 μg of cell lysate was incubated with 25 μl of streptavidin agarose beads in RIPA buffer for 3 hours at 4°C with rotation. Beads were washed with 10 volumes of RIPA buffer, resuspended in sample buffer (LDS), and heated at 75°C for 10 minutes. The resulting samples were analyzed by sodium dodecyl sulfate-polyacrylamide gel electrophoresis (SDS–PAGE), followed by western blot analysis described previously (18).

#### Western Blot Analysis

For western blot analysis, 15 μg of total cell lysate was used for total protein detection. The following antibodies were used:

- Anti-c-Myc Antibody (9E10; monoclonal IgG; dilution 1:1000), Thermo Fisher Scientific (St. Louis, MO).
- Anti-HSP70 Monoclonal Antibody (Mouse IgG; clone C92F3A-5; dilution 1:1000), Enzo Life Sciences, Farmingdale, NY, USA.
- THE™ Beta Actin Antibody (Mouse mAb; A00702; dilution 1:1000), GenScript, Piscataway, NJ, USA.
- Na⁺/K⁺ ATPase α (ATP1A1) Antibody RabMAb® (EP1845Y; 2047-1; dilution 1:1000), Thermo Fisher Scientific (St. Louis, MO).

Blots were incubated with primary antibodies overnight (∼16 hours) at 4°C with constant rotation. Total protein staining was performed using Pierce™ Reversible Protein Stain (Thermo Scientific Pierce, Rockford, IL, USA).

#### Signal Detection

Western blot signals were visualized using the Omega Lum™ C Imaging System (Aplegen, San Francisco, CA).

### Mass Spectrometry and Lipidome Analysis

#### Sample Preparation

Six biological replicates (different cell batches) of four million HeLa cells each were prepared to identify total lipid profiles before and after heat shock. Samples were collected under three conditions: control (37°C), immediately after heat shock (R0), and after an 8-hour recovery period following heat shock (R8).

#### Lipid Extraction and Analysis

Samples were analyzed by the UC Davis West Coast Metabolomics Center using biphasic lipid extraction, followed by data acquisition on a C18-based hybrid bridged column with a ternary water/acetonitrile/isopropanol gradient and a ThermoFisher Scientific Q-Exactive HF mass spectrometer with electrospray ionization. MS/MS data were acquired in data-dependent mode, and accurate masses were normalized by constant reference ion infusion. MS-DIAL vs. 4.90 was used for data processing and lipid annotations (47,48). Lipids were annotated in identity search mode with precursor mass errors < 10 mDa, retention time matching to the corresponding lipid classes and MS/MS matching. Data were sum-normalized: each lipid was represented as a fraction of total lipids in the sample and scaled by the median intensity of all lipids within the treatment group (49–51). Data are shown in Supplemental Table 1. Heatmap analysis (MetaboAnalyst (52)) was performed to simultaneously visualize lipidomic changes in response to experimental conditions and cluster similar samples based on their lipid profiles. Total lipid abundance was visualized using boxplot graphs (53).

### Statistical Tests

Statistical significance was assessed using one-way ANOVA (Analysis of Variance) followed by post-hoc Tukey HSD (Honestly Significant Difference) and Bonferroni tests. A *P* value < 0.05 was considered statistically significant. Boxplots were generated using the BoxPlotR application (http://shiny.chemgrid.org/boxplotr/) (53).

## Results

### HSPA1A Plasma Membrane Localization Peaks Following Heat Shock

To investigate the hypothesis that phosphatidylserine (PS) regulates the translocation of HSPA1A to the plasma membrane (PM) during cellular stress, we first quantified the timing and extent of HSPA1A’s PM localization following heat shock (Fig. 1). Using confocal microscopy, we observed that HSPA1A levels at the PM increased significantly after heat shock, peaking around 8 hours post-stress, and subsequently returning to control levels by 24 hours (Fig. 1A). Additionally, cell-surface biotinylation and subsequent western blot analysis confirmed a marked increase in PM-localized HSPA1A at 8 hours (Fig. 1B and Supplemental Fig. 1), aligning with our imaging data. These results indicate a dynamic and temporally regulated localization pattern of HSPA1A at the PM, corresponding with the recovery phase of the heat shock response.

**Figure 1.**
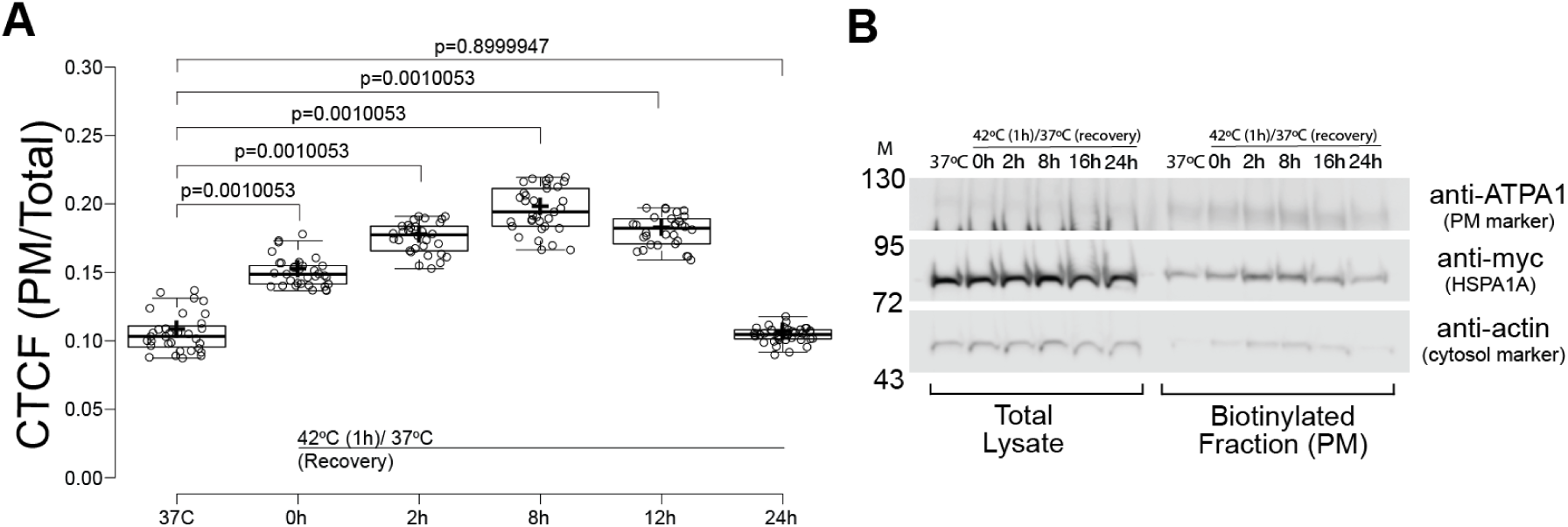
HSPA1A’s plasma membrane (PM) localization significantly increases during recovery from mild heat shock. (A) Quantification of confocal imaging data showing the corrected total cell fluorescence (CTCF) ratio of HSPA1A fluorescence at the PM to the rest of the cell. Localization was analyzed in cells under control conditions (37°C) or after heat shock (1 h at 42°C), followed by recovery at 37°C for 0–24 h. Data represent three independent experiments, with individual cells shown as open circles (total N=30). The center lines show medians; box limits indicate the 25th and 75th percentiles; whiskers extend 1.5 times the interquartile range; crosses represent sample means. Statistical significance was determined using one-way ANOVA (Analysis of Variance) followed by a post-hoc Tukey HSD (Honestly Significant Difference) test. (B) Cell surface biotinylation confirms that HSPA1A’s PM association significantly increases in response to heat shock, mirroring the confocal imaging results. Representative cropped. Western blots show total and biotinylated fractions from HEK293 cell lysates expressing HSPA1A-myc (complete blots in Supplemental Figure 1). Actin and ATP1A1 were used as cytosolic and membrane controls, respectively. M: molecular size marker (Fisher BioReagents™ EZ-Run™ Prestained Rec Protein Ladder; approximate sizes indicated on the left).

These findings support the notion that HSPA1A’s PM translocation is not merely a passive diffusion process but is potentially mediated by specific lipid interactions, particularly with PS. To further explore the role of PS in this translocation process, we next assessed how manipulating PS levels would affect HSPA1A’s PM localization. Using pharmacological and genetic approaches to inhibit PS synthesis, we sought to determine whether PS increase following heat shock triggers HSPA1A’s movement to the PM.

### Phosphatidylserine (PS) Increase is Required for HSPA1A Membrane Localization

We first quantified PS levels to test our hypothesis that PS is essential for HSPA1A’s translocation to the plasma membrane (PM) in response to heat shock. Using confocal microscopy to monitor the PS-specific biosensor Lact-C2 and lipidomics analysis, we observed that PS levels at the PM significantly increased immediately following heat shock (0h) and then dropped during recovery (8h) (Figs. 2-3 and Supplemental Fig. 2). This temporal pattern of PS elevation aligns closely with the observed HSPA1A translocation, supporting that PS accumulation at the PM may be a critical signal for HSPA1A recruitment.

**Figure 2.**
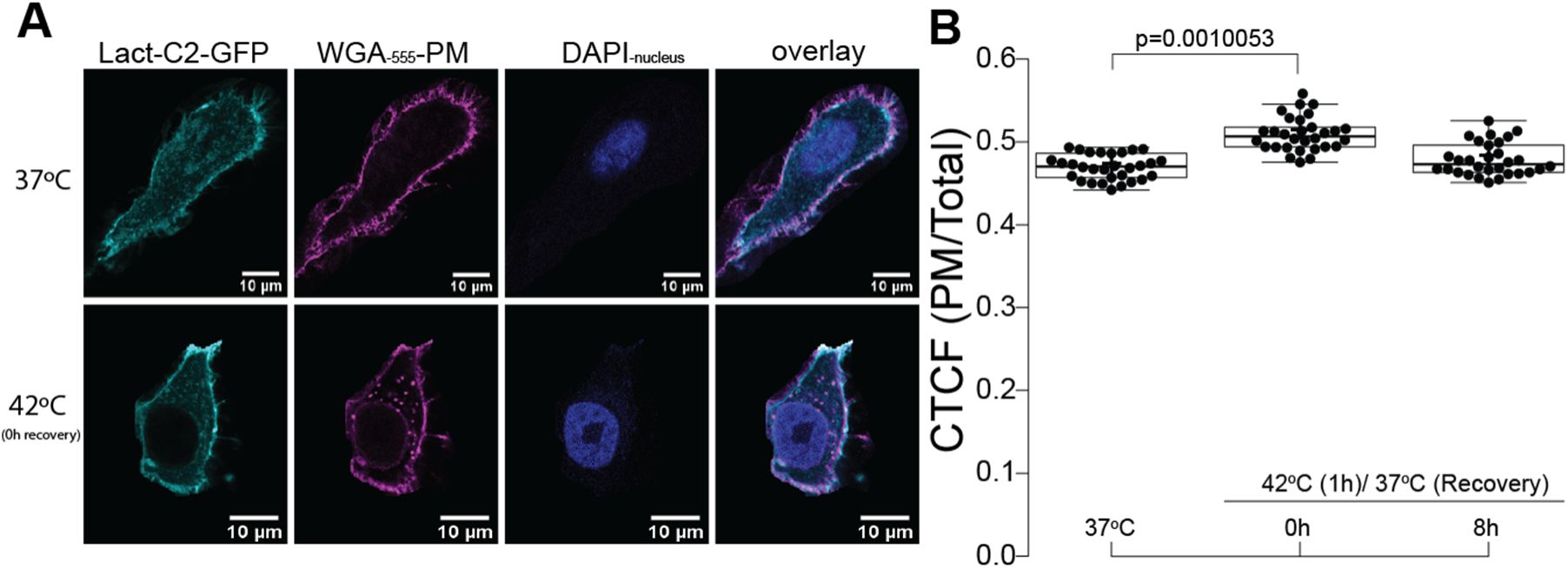
Heat shock increases phosphatidylserine (PS) levels at the plasma membrane (PM) as shown by Lact-C2’s (PS-biosensor) localization. (A) Representative confocal images of HeLa cells expressing GFP-Lact-C2, a PS biosensor, stained with WGA-FA555 (PM stain) and DAPI (nucleus stain). Lact-C2 localization was analyzed under control conditions (37°C; top row) and heat shock (1 h at 42°C; no recovery; bottom row). Scale bar = 10 μm. (B) Quantification of the corrected total cell fluorescence (CTCF) ratio of Lact-C2 fluorescence at the PM to the rest of the cell under control and heat-shocked conditions. Data represent three independent experiments, with total individual cell counts (N = 30) shown at the bottom of the graph. Center lines show medians; box limits indicate the 25th and 75th percentiles; whiskers extend 1.5 times the interquartile range; crosses represent sample means. Statistical significance was determined using one-way ANOVA (Analysis of Variance) followed by post-hoc Tukey HSD (Honestly Significant Difference) test.

**Figure 3.**
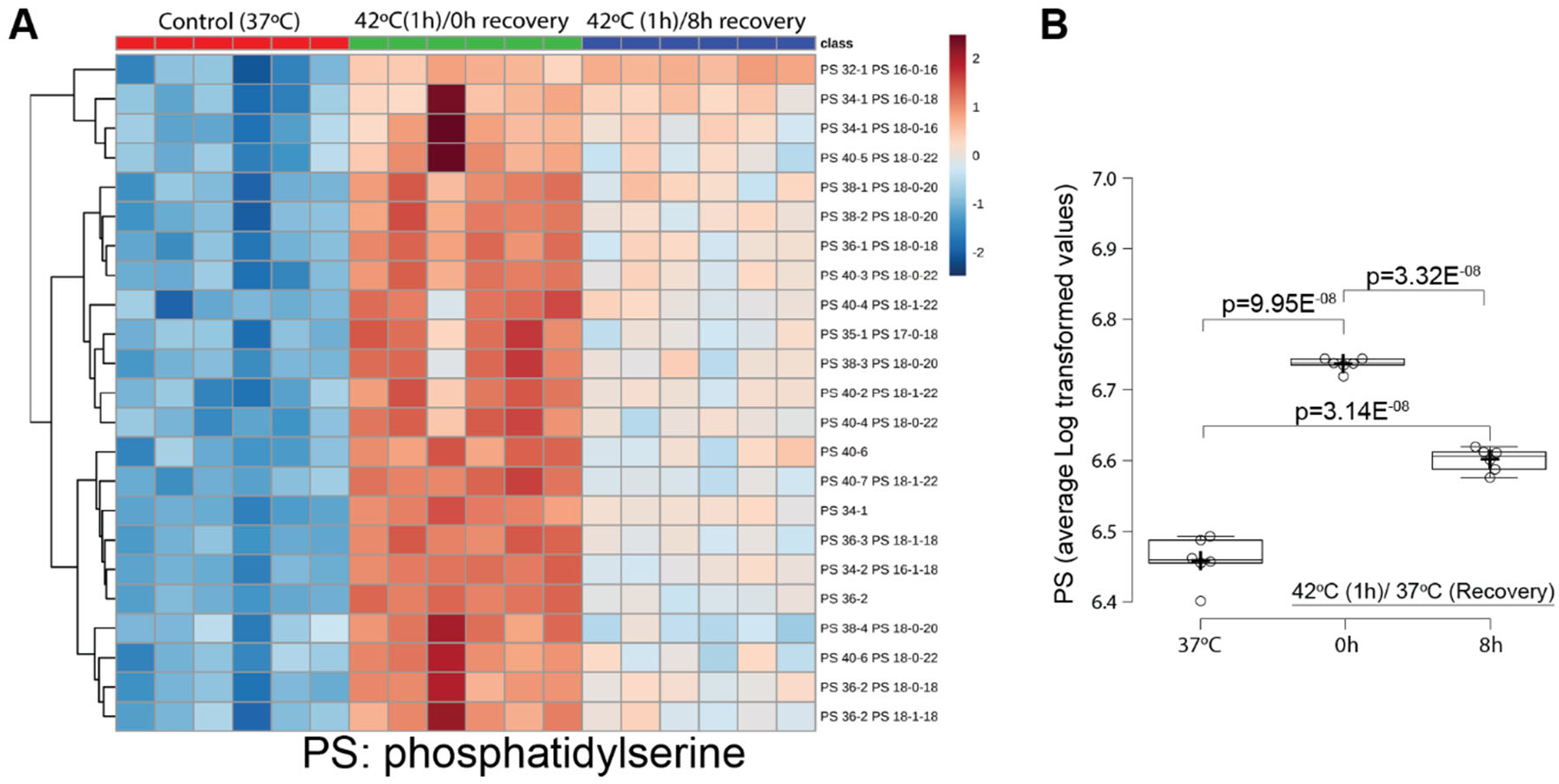
Total cellular phosphatidylserine (PS) increases immediately following heat shock. (A) Heatmap visualization of PS species composition in HeLa cells under control conditions (37°C), after heat shock (1 h at 42°C; 0 h recovery) and following 8 h of recovery at 37°C. Lipid composition was calculated as the percentage of each PS subclass within the total lipidome and log-transformed for visualization. Each column represents one biological replicate. The heatmap was generated using MetaboAnalyst (52). (B) Boxplot showing the total PS levels across six biological replicates, calculated from log-transformed data of all PS species in each replicate. Center lines indicate medians; box limits represent the 25th and 75th percentiles; whiskers extend 1.5 times the interquartile range; crosses denote sample means. Statistical significance was determined using one-way ANOVA (Analysis of Variance) followed by post-hoc Tukey HSD (Honestly Significant Difference) and Bonferroni tests.

We applied fendiline to inhibit PS synthesis to investigate whether PS is required for HSPA1A’s PM localization. Confocal imaging with the Lact-C2 biosensor confirmed that fendiline effectively reduced PM-associated PS levels. Correspondingly, HSPA1A levels at the PM decreased significantly in fendiline-treated cells, as shown by imaging data (Fig. 4 and Supplemental Fig. 3). These results indicate that the increase in PS following heat shock is necessary for HSPA1A translocation to the PM.

**Figure 4.**
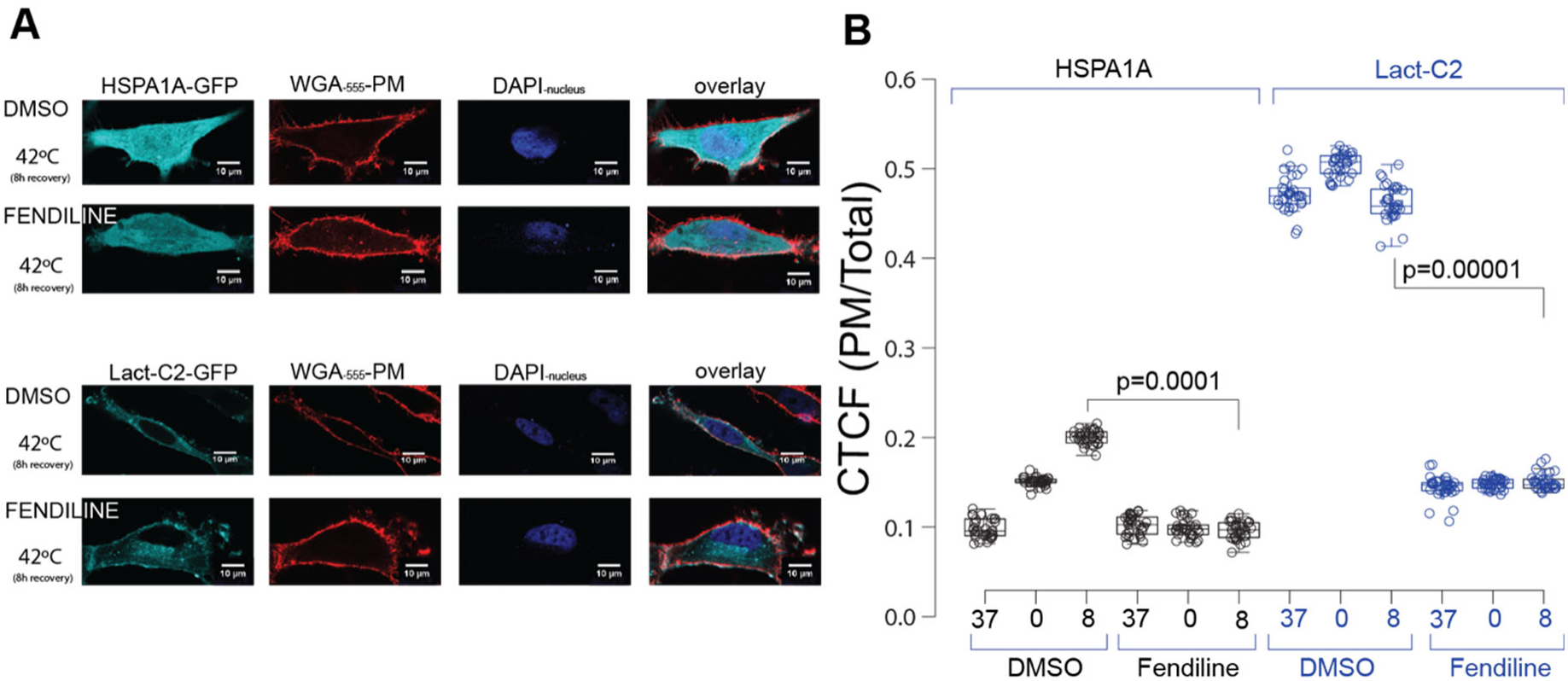
Fendiline treatment significantly decreases HSPA1A’s plasma membrane (PM) localization after heat shock. (A) Representative images of HeLa cells expressing GFP-HSPA1A (top panels) or GFP-Lact-C2 (bottom panels). Cells were either untreated (DMSO; top rows in each panel) or treated with 10 µM fendiline for 48 hours (bottom rows in each panel). Images show cells after heat shock (1 h at 42°C) followed by 8 hours of recovery (additional images can be found in Supplemental Figure 3). PM was stained with WGA-FA555, and nuclei were stained with DAPI (second and third images from the left in each panel). Scale bar = 10 µm. (B) Quantification of the corrected total cell fluorescence (CTCF) as a ratio of total fluorescence at the PM to fluorescence in the rest of the cell under control conditions and after heat shock [directly after heat shock (0 hours of recovery) and 8 hours of recovery], with or without fendiline treatment. The experiment was performed three times, with a total of 30 cells per condition (shown as open circles). Box limits indicate the 25th and 75th percentiles, whiskers extend 1.5 times the interquartile range, and crosses represent sample means. Statistical significance was determined using one-way ANOVA (Analysis of Variance) followed by post-hoc Tukey HSD (Honestly Significant Difference) and Bonferroni tests.

Further confirming the role of PS, we used RNA interference (RNAi) to target key enzymes in PS synthesis, phosphatidylserine synthase 1 and 2 (PSS1, PSS2). Knockdown of PSS1 and PSS2 (via RNAi against PTDSS1 and PTDSS2 genes) led to reduced Lact-C2 (PS biosensor) and HSPA1A localization at the PM following heat shock (Fig. 5 and Supplemental Fig. 4), consistent with our findings using fendiline. These data strongly suggest that PS increase is crucial for HSPA1A’s PM recruitment under heat shock.

**Figure 5.**
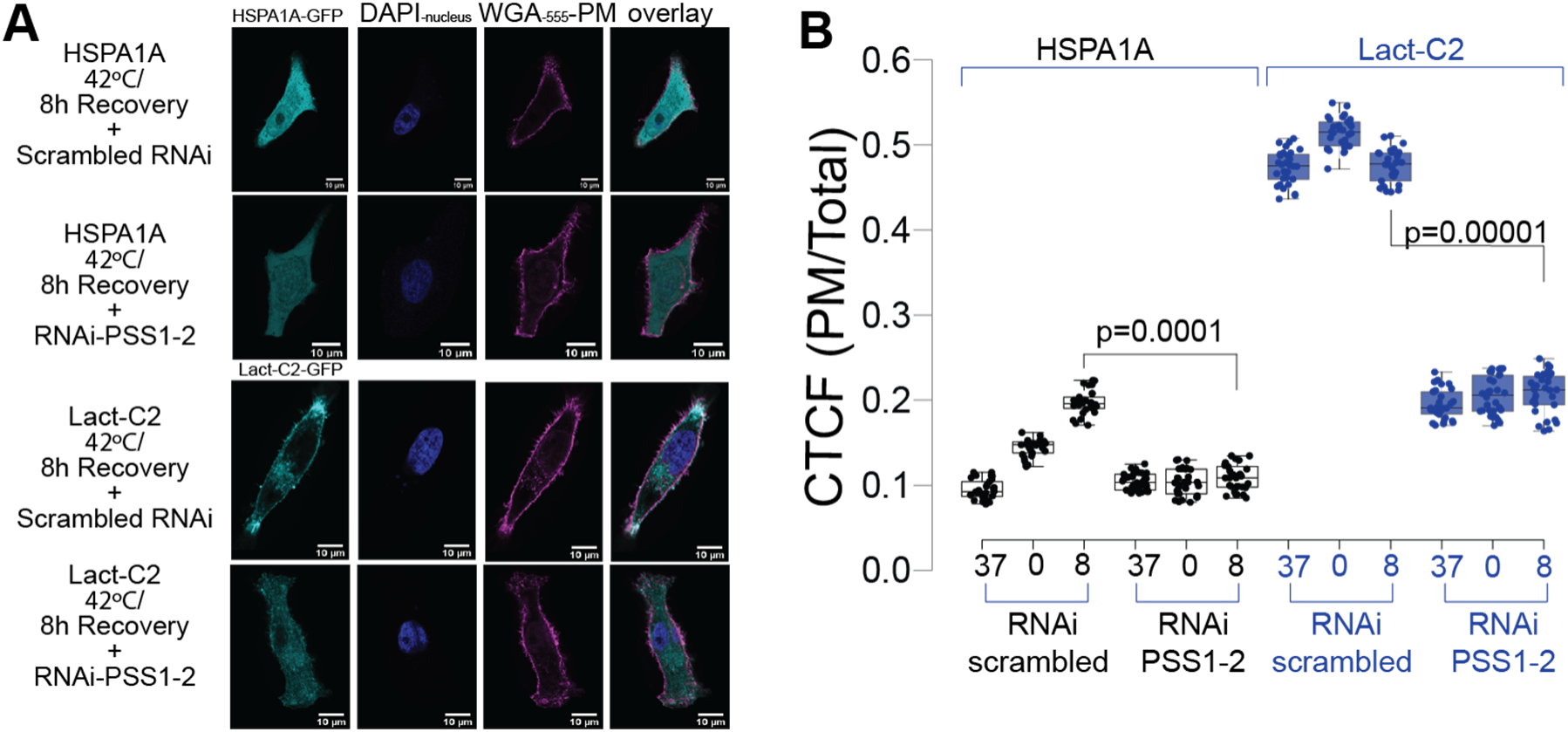
HSPA1A plasma membrane (PM) localization significantly decreases when PTDSS1 and PTDSS2 genes are silenced using RNAi. (A) Representative images of HeLa cells expressing HSPA1A-GFP (top panels) and GFP-Lact-C2 (bottom panels). Cells were transfected with either scrambled RNAi (top rows in each panel) or RNAi targeting the PS synthesis genes *PTDSS1* and *PTDSS2* (bottom rows in each panel). Images show cells after heat shock (1 h at 42°C) followed by 8 hours of recovery. PM was stained with WGA-FA555, and nuclei were stained with DAPI (second and third images from the left in each panel). Scale bar = 10 µm. (B) Quantification of corrected total cell fluorescence (CTCF) as the ratio of total GFP fluorescence at the PM to the rest of the cell. Measurements were performed under control conditions and after heat shock [immediately after heat shock (0 hours of recovery) and following 8 hours of recovery] with simultaneous silencing of *PTDSS1* and *PTDSS2*. The experiment was repeated three times, with a total of 30 cells per condition (shown as closed circles). Box limits indicate the 25th and 75th percentiles, whiskers extend 1.5 times the interquartile range, and crosses represent sample means. Statistical significance was determined using one-way ANOVA (Analysis of Variance) followed by post-hoc Tukey HSD (Honestly Significant Difference) and Bonferroni tests. Results from additional conditions are shown in Supplemental Figure 4.

To explore whether additional lipid modifications might influence HSPA1A’s membrane localization, we next examined the effects of modulating levels of other lipids on HSPA1A’s PM localization.

### Specificity of HSPA1A Localization to PS Among Heat-Shock-Induced Lipids

To evaluate whether HSPA1A’s PM localization is uniquely dependent on phosphatidylserine (PS) or if other heat shock-induced lipids also contribute, we examined the effects of additional lipids that increase following heat shock.

Lipidomic analysis showed cholesterol levels rose substantially post-heat shock, with total cellular cholesterol levels elevated, while PM-specific cholesterol showed no significant change (Fig. 6).

**Figure 6.**
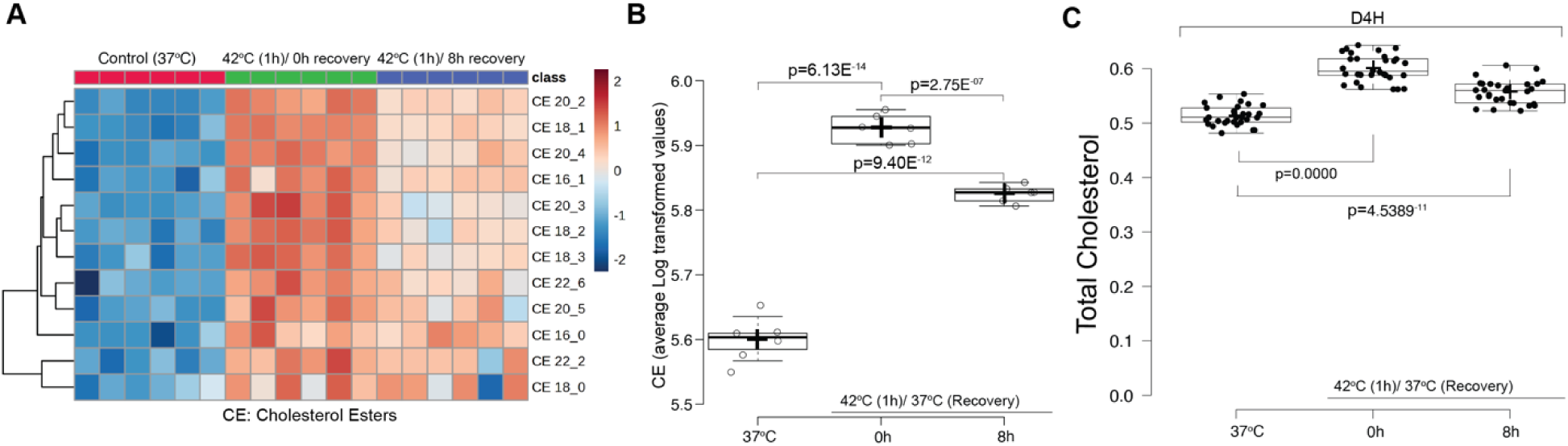
Total cellular cholesterol esters (CE) increase after heat shock. (A) Heatmap visualization of CE species composition in HeLa cells under control conditions (37°C), after heat shock (1 h at 42°C, 0 h recovery), and following 8 h of recovery at 37°C. Lipid composition was calculated as the percentage of each CE subclass within the total lipidome and log-transformed for visualization. Each column represents one biological replicate. The heatmap was generated using MetaboAnalyst (52). (B) Boxplot showing the total CE levels across six biological replicates, calculated from log-transformed data of all CE species in each replicate. Center lines indicate medians; box limits represent the 25th and 75th percentiles; whiskers extend 1.5 times the interquartile range; crosses denote sample means. (C) Quantification of corrected total cell fluorescence (CTCF) from confocal images, representing total cholesterol (total D4H fluorescence signal) per cell under control conditions (37°C) and after heat shock (0 h and 8 h recovery). The experiment was performed three times, with a total of 30 cells per condition (shown as closed circles). Box limits indicate the 25th and 75th percentiles; whiskers extend 1.5 times the interquartile range; crosses represent sample means. For both B and C, statistical significance was determined using one-way ANOVA (Analysis of Variance) followed by post-hoc Tukey HSD (Honestly Significant Difference) and Bonferroni tests.

Using the cholesterol-binding probe D4H, which masks cholesterol at the PM, we confirmed that PM cholesterol masking had minimal impact on HSPA1A localization (Fig. 7). This finding was reinforced by using methyl-beta-cyclodextrin (MβCD) to deplete PM cholesterol, which showed minimal effects on HSPA1A’s PM presence, suggesting cholesterol does not significantly influence HSPA1A localization after heat shock (Fig. 8).

**Figure 7.**
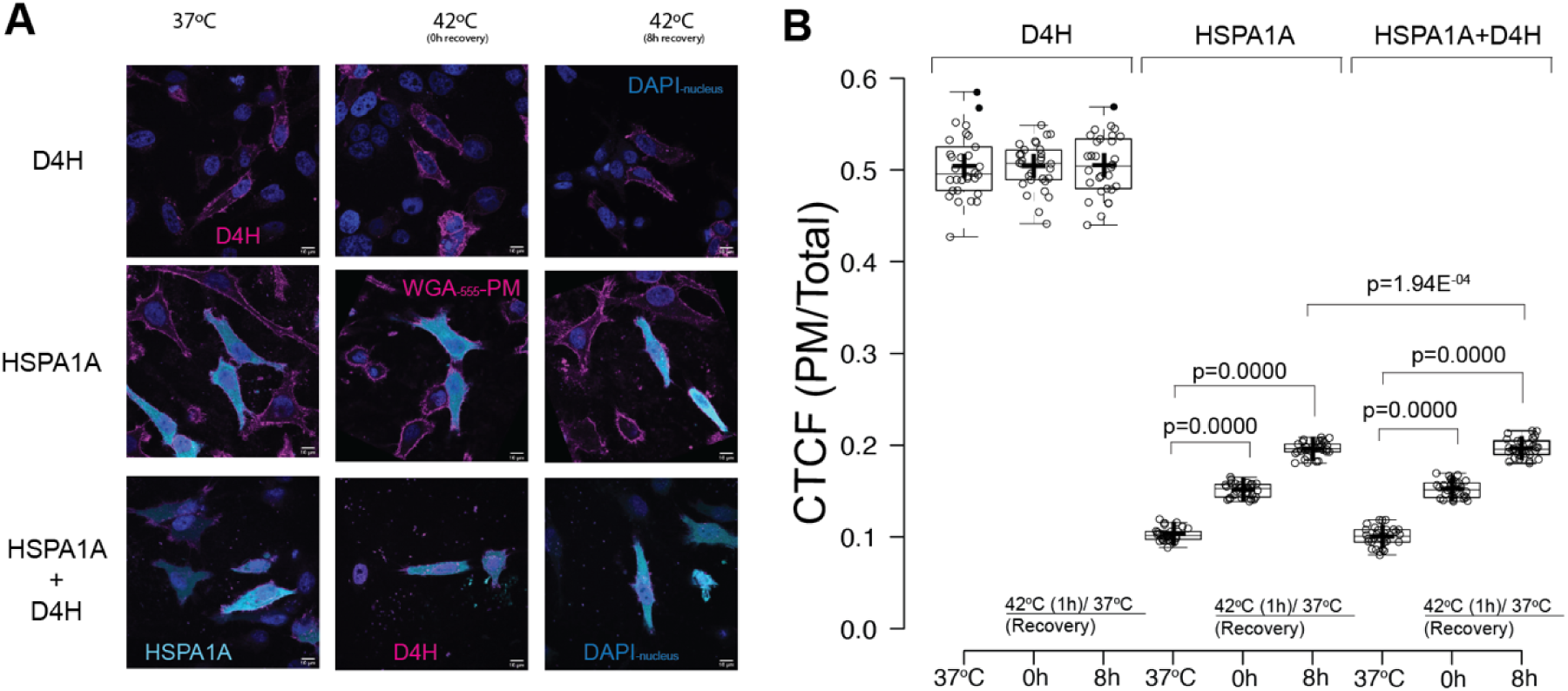
Plasma membrane (PM) cholesterol masking using the cholesterol biosensor D4H results in minimal change in HSPA1A’s PM localization after heat shock. (A) Representative images of HeLa cells expressing the cholesterol biosensor D4H-RFP (top panels, magenta), GFP-HSPA1A (middle panels, cyan), or both D4H-RFP and GFP-HSPA1A (bottom panels). Images show cells under control conditions (37°C), immediately after heat shock (1 h at 42°C, 0 hours of recovery), or following 8 hours of recovery at 37°C (8h). The PM was stained with WGA-FA555 (magenta; middle panels), and nuclei were stained with DAPI (blue; all panels). Scale bar = 10 µm. (B) Quantification of corrected total cell fluorescence (CTCF) as the ratio of total fluorescence at the PM to fluorescence in the rest of the cell under control conditions and after heat shock (0h and 8h). The experiment was performed three times, with a total of 30 cells per condition (shown as open circles). Box limits indicate the 25th and 75th percentiles, whiskers extend 1.5 times the interquartile range, and crosses represent sample means. Statistical significance was determined using one-way ANOVA (Analysis of Variance) followed by post-hoc Tukey HSD (Honestly Significant Difference) and Bonferroni tests.

**Figure 8.**
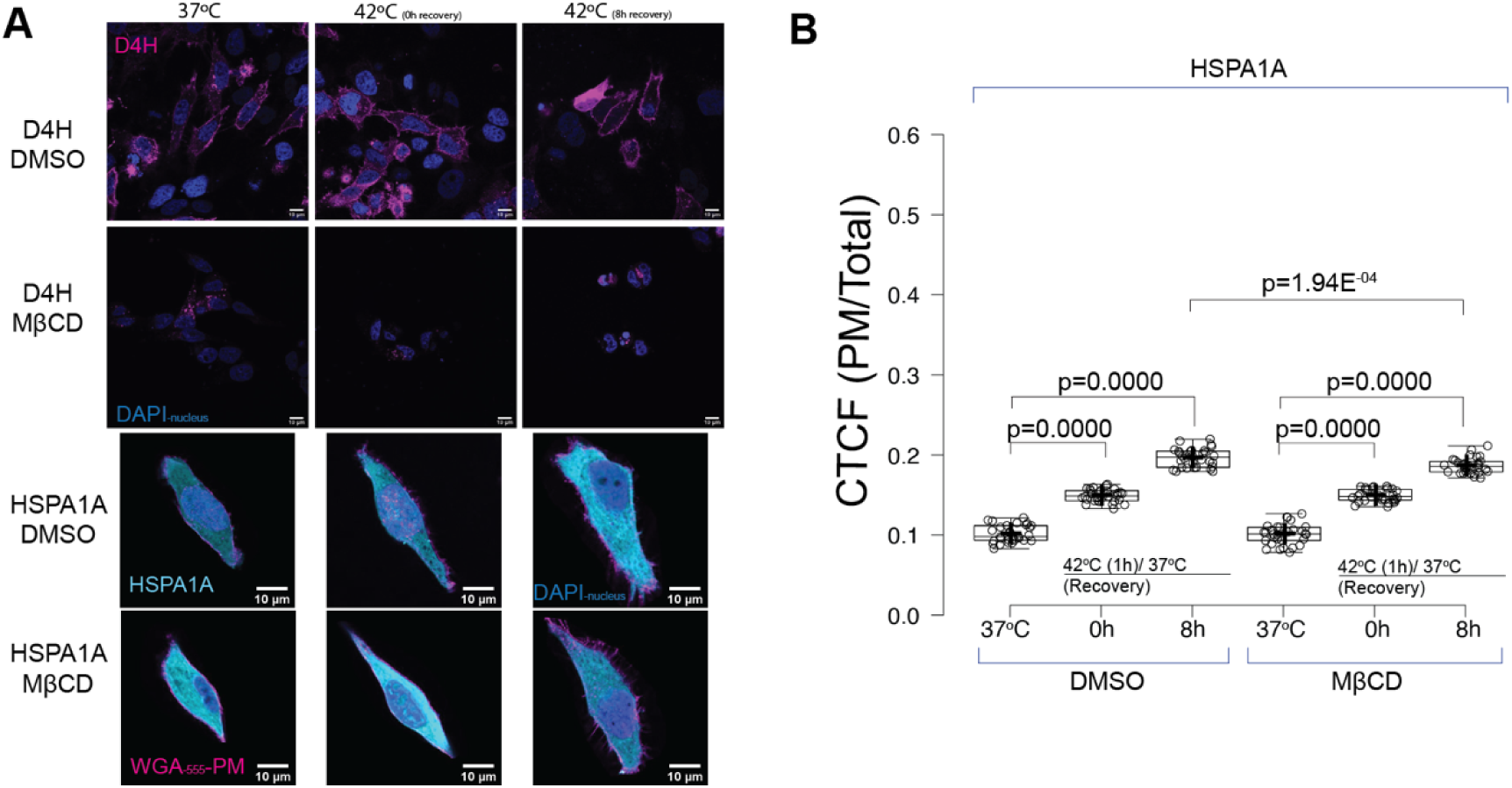
Plasma membrane (PM) cholesterol depletion using methyl-β-cyclodextrin (MβCD) results in minimal change in HSPA1A’s PM localization after heat shock. (A) Representative images of HeLa cells expressing the cholesterol biosensor D4H-RFP (top panels, magenta) or GFP-HSPA1A (bottom panels, cyan). Cells were either untreated (DMSO; top rows in each panel) or treated with 5 mM MβCD for 9 hours (bottom rows in each panel). Images show cells under control conditions (37°C), immediately after heat shock (1 h at 42°C, 0 hours of recovery), or following 8 hours of recovery at 37°C (8h). PM was stained with WGA-FA555 (magenta), and nuclei were stained with DAPI (blue). Scale bar = 10 µm. (B) Quantification of corrected total cell fluorescence (CTCF) as the ratio of total fluorescence at the PM to fluorescence in the rest of the cell under control conditions and after heat shock (0h and 8h), with or without MβCD treatment. The experiment was performed three times, with a total of 30 cells per condition (shown as open circles). Box limits indicate the 25th and 75th percentiles, whiskers extend 1.5 times the interquartile range, and crosses represent sample means. Statistical significance was determined using one-way ANOVA (Analysis of Variance) followed by post-hoc Tukey HSD (Honestly Significant Difference) and Bonferroni tests.

Next, we investigated fatty acids, another lipid category upregulated by heat shock (Fig. 9). Suppression of fatty acid synthase (FASN), the enzyme responsible for fatty acid synthesis, slightly decreased HSPA1A’s PM localization compared to untreated controls (Fig. 10). However, the effect remained minor relative to the impact of PS inhibition. These data indicate that HSPA1A’s PM translocation is specifically sensitive to PS, as modulating other heat shock-induced lipids like cholesterol and fatty acids produced only minimal changes in HSPA1A localization.

**Figure 9.**
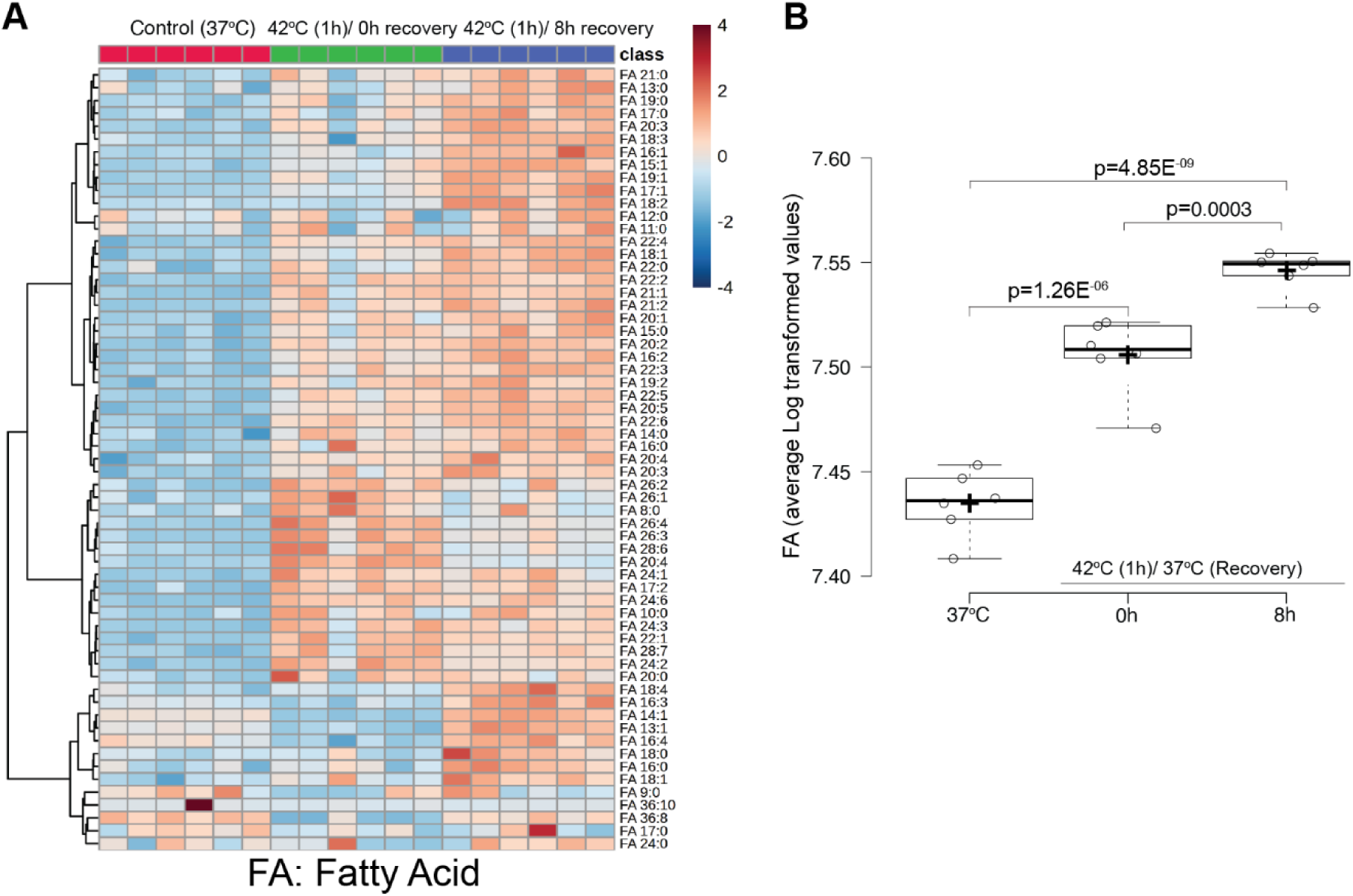
Total cellular fatty acids (FA) increase after heat shock. Heatmap visualization of FA species composition in HeLa cells under control conditions (37°C), immediately after heat shock (1 h at 42°C, 0 h recovery), and following 8 h of recovery at 37°C. Lipid composition was calculated as the percentage of each FA subclass within the total lipidome and log-transformed for visualization. Each column represents one biological replicate. The heatmap was generated using MetaboAnalyst (52). (B) Boxplot showing the total FA levels across six biological replicates, calculated from log-transformed data of all FA species in each replicate. Center lines indicate medians; box limits represent the 25th and 75th percentiles; whiskers extend 1.5 times the interquartile range; crosses denote sample means. Statistical significance was determined using one-way ANOVA (Analysis of Variance) followed by post-hoc Tukey HSD (Honestly Significant Difference) and Bonferroni tests.

**Figure 10.**
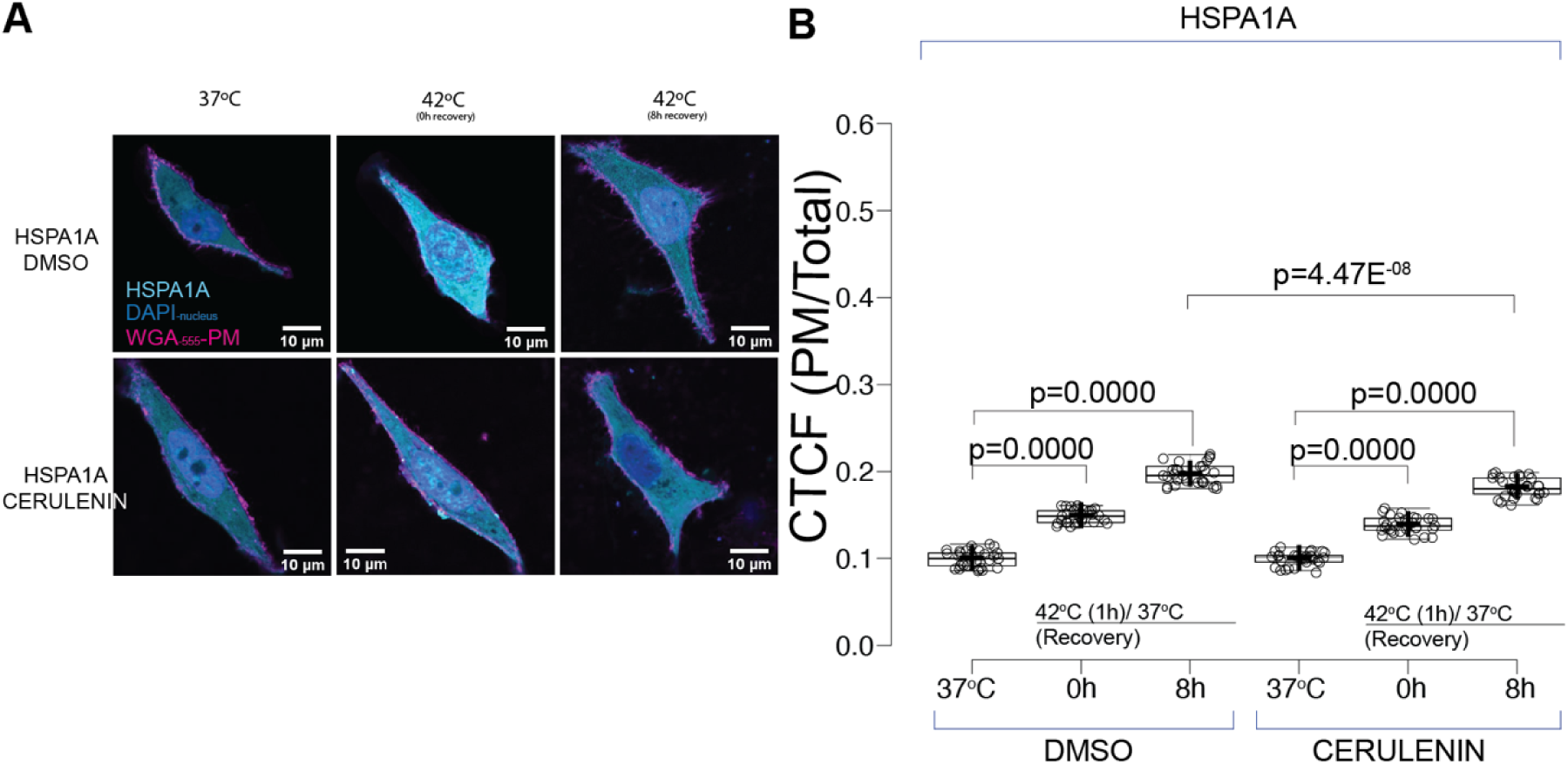
Inhibition of fatty acid synthesis using cerulenin, a fatty acid synthase (FASN) inhibitor, results in minimal change in HSPA1A’s PM localization after heat shock. (A) Representative images of HeLa cells expressing GFP-HSPA1A (cyan). Cells were either untreated (DMSO; top row) or treated with 10 µg/ml cerulenin for 9 hours (bottom row). Images depict cells under control conditions (37°C), immediately after heat shock (1 h at 42°C, 0 hours of recovery), or following 8 hours of recovery at 37°C (8h). The PM was stained with WGA-FA555 (magenta), and nuclei were stained with DAPI (blue). Scale bar = 10 µm. (B) Quantification of corrected total cell fluorescence (CTCF) as the ratio of total fluorescence at the PM to fluorescence in the rest of the cell under control conditions and after heat shock (0h and 8h), with or without cerulenin treatment. The experiment was performed three times, with a total of 30 cells per condition (shown as open circles). Box limits indicate the 25th and 75th percentiles, whiskers extend 1.5 times the interquartile range, and crosses represent sample means. Statistical significance was determined using one-way ANOVA (Analysis of Variance) followed by post-hoc Tukey HSD (Honestly Significant Difference) and Bonferroni tests.

Given the specificity of PS’s role in driving HSPA1A localization, our next objective was to identify which properties of PS—such as its acyl chain saturation—might further influence this process.

### PS Saturation and Chain Length Do Not Affect HSPA1A Localization

To further refine our understanding of how PS regulates HSPA1A’s PM localization, we examined whether the specific molecular characteristics of PS—namely, saturation and chain length—might influence this process. Given that cancer cells often exhibit altered membrane composition, including lipid saturation, and that HSPA1A embedding into artificial membranes correlates with PS saturation levels, we theorized that heat shock-induced changes in PS species could impact HSPA1A’s translocation to the PM.

Our lipidomic analysis revealed that following heat shock, the carbon chain length and saturation level of PS increased, indicating substantial structural modifications to PS at the PM (Fig. 11).

**Figure 11.**
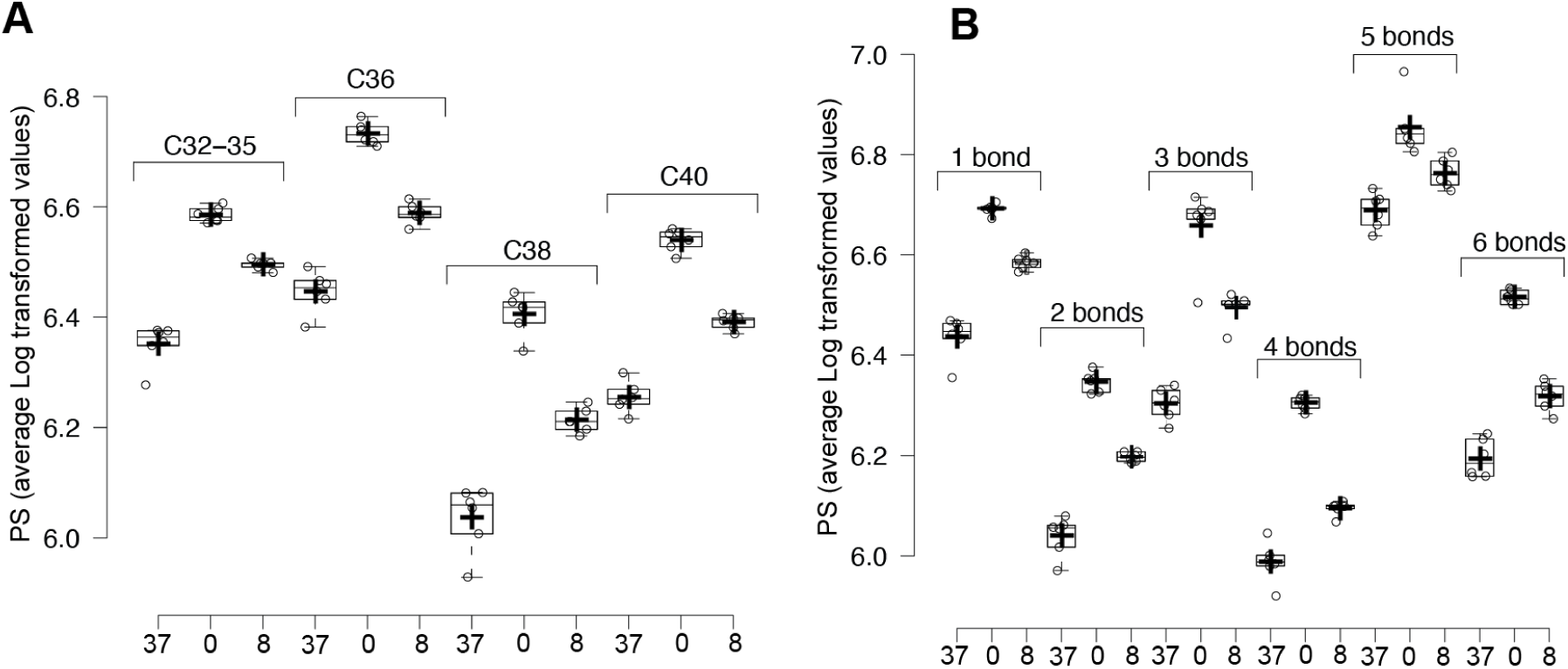
Acyl-chain length and saturation levels of PS show a sharp increase immediately following heat shock. (A) Boxplots showing the averages of PS species categorized by carbon chain length. Each boxplot represents the average, calculated from the log-transformed data of all PS species in the respective category, across six biological replicates under control (37°C) or heat shock conditions (1 h at 42°C, 0 h recovery). (B) Boxplots showing the averages of PS species categorized by the number of double bonds. Each boxplot represents the average, calculated from the log-transformed data of all PS species in the respective category, across six biological replicates under the same conditions. For both panels, center lines indicate medians; box limits represent the 25th and 75th percentiles; whiskers extend 1.5 times the interquartile range, and crosses denote sample means.

To determine if these specific alterations were necessary for HSPA1A localization, we employed pharmacological inhibitors (CP and SC) to block desaturase activity, thereby stabilizing PS saturation levels.

This desaturase inhibition caused minimal changes in HSPA1A’s PM localization compared to controls, suggesting that variations in PS acyl chains alone do not significantly affect HSPA1A’s membrane localization (Fig. 12). Therefore, while PS saturation and chain length are modified post-heat shock, these specific characteristics do not appear to govern HSPA1A’s PM translocation. Instead, the total increase in PS levels is the key determinant of HSPA1A’s localization.

**Figure 12.**
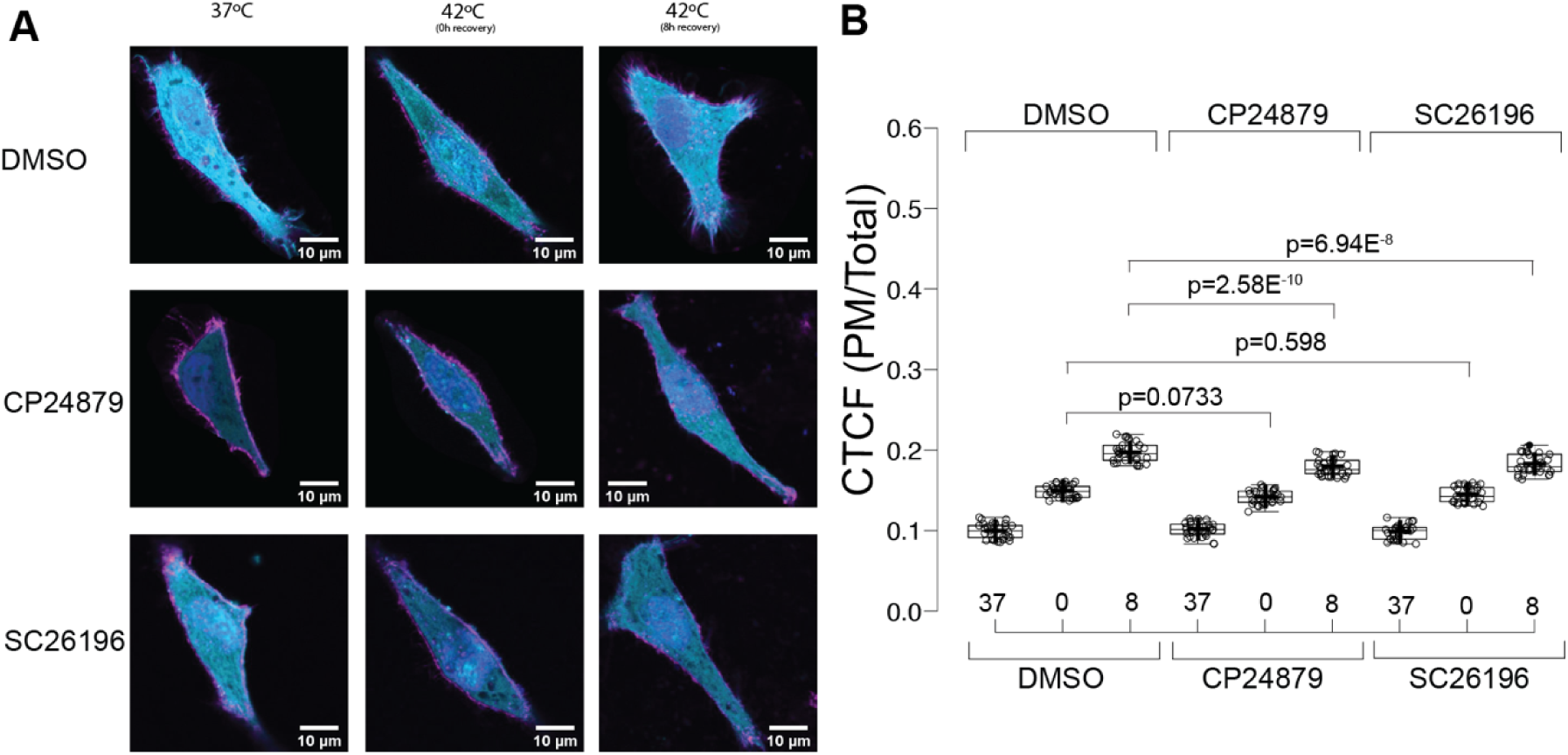
Desaturase inhibitors have minimal effect on HSPA1A’s PM localization after heat shock. (A) Representative images of HeLa cells expressing GFP-HSPA1A (cyan). Cells were either untreated (DMSO; top row) or treated with 5.5 µM CP-24879 (middle row) or 100 µM SC 26196 (bottom row) for 48 hours. Images show cells under control conditions (37°C), immediately after heat shock (1 h at 42°C, 0 h recovery), or following 8 hours of recovery at 37°C (8h). The PM was stained with WGA-FA555 (magenta), and nuclei were stained with DAPI (blue). Scale bar = 10 µm. (B) Quantification of corrected total cell fluorescence (CTCF) as the ratio of total GFP fluorescence at the PM to fluorescence in the rest of the cell, under control conditions and after heat shock (0h and 8h), with or without desaturase inhibitor treatments. The experiment was repeated three times, with a total of 30 cells per condition (shown as open circles). Box limits represent the 25th and 75th percentiles, whiskers extend 1.5 times the interquartile range, and crosses denote sample means. Statistical significance was determined using one-way ANOVA (Analysis of Variance) followed by post-hoc Tukey HSD (Honestly Significant Difference) and Bonferroni tests.

Collectively, our findings support the hypothesis that elevated PS levels after heat shock are both necessary and sufficient to drive HSPA1A’s movement to the PM. The consistent timing, PS dependency, and sufficiency of PS increase establish a direct, causal link between PS upregulation and HSPA1A translocation.

## Discussion

Our findings reveal that heat shock triggers specific lipid changes, including increased PS levels at the plasma membrane. This observation confirms that heat stress induces significant lipidomic alterations across various organisms (26,54,55). The observed rise in PS underscores its role as a critical early signal in the heat shock response, where lipid composition changes and membrane reorganization serve as early biochemical signals for cellular stress.

This lipidome shift highlights membrane composition’s dual role: providing structural support and actively participating in stress signaling pathways. Heat-induced membrane reorganization, including lipid rafts and fluidity changes, has been shown to impact protein function and cellular signaling (56,57). Similarly, PS exposure during stress is implicated in diverse signaling processes (58,59), supporting its regulatory significance. Alterations in membrane fluidity also activate heat shock factor 1 (HSF1), linking lipid changes to transcriptional regulation during stress (60,61). These findings reinforce the idea that membrane lipids like PS are integral components of the cellular stress response machinery.

Our results specifically reveal that PS elevation acts as a lipid-specific signal orchestrating the mobilization of HSPA1A to the plasma membrane. This finding aligns with the growing recognition of lipids as crucial signaling molecules in stress responses (57,62). By identifying PS as a lipid altered by heat shock, we provide evidence for a molecular mechanism by which cells adapt to stress, engaging specific lipid molecules to mobilize protective chaperones like HSPA1A (14,19,63). Unlike other lipids, such as cholesterol and fatty acids, which showed minimal effects on HSPA1A localization, PS uniquely governs HSPA1A’s translocation. This specificity underscores the role of lipid signals in controlling chaperone relocalization during cellular stress.

The PS-driven translocation of HSPA1A (14) underscores lipid alterations as integral to stress signaling, where changes in membrane composition activate molecular chaperones’ downstream functions. Research has shown that membrane lipid composition modulates heat shock protein function and localization (55,60), and our findings extend this knowledge by identifying PS as a critical lipid partner. PS-dependent recruitment of proteins, reminiscent of its role in apoptosis signaling (59), now emerges as a novel mechanism regulating HSPA1A during heat shock responses.

This study also broadens our understanding of the lipid-chaperone interface. By revealing that HSPA1A’s PM localization is selectively dependent on PS, we confirm the specificity of lipid-protein interactions in membrane environments (57,62). This insight suggests that lipid identity fine-tunes chaperone activity (3,14,31,63) according to cellular needs, adding depth to our understanding of how membrane lipids regulate protein function during stress. The broader implications of PS regulation extend to cancer biology, where lipid remodeling is common.

Our data add to growing evidence that altered lipid compositions regulate cellular stress pathways, with implications for cancer. PS levels at the PM, which increase during heat shock and in many cancers, appear to initiate HSPA1A membrane localization. This concept links the chaperone’s activity to cellular processes such as proteostasis and vesicular trafficking, supporting studies that show lipid alterations modulate heat shock protein function and localization (3,55,60). These PS-dependent mechanisms likely underpin HSPA1A’s roles in cancer cell survival and therapeutic resistance. Heat shock proteins, including HSPA1A, are well-established contributors to cancer progression and drug resistance (22,64,65), and lipid metabolism reprogramming in cancer cells often aids their survival (23,66).

Given the association between HSPA1A PM localization and therapeutic resistance in cancer (7,67–70), our study suggests PS modulation as a potential therapeutic target. Interventions manipulating PS levels or lipid metabolism could inhibit HSPA1A’s stress-protective functions. Pharmacological agents targeting lipid metabolism have shown promise in sensitizing cancer cells to therapy (66,71), and our findings open the door to similar strategies aimed at preventing HSPA1A membrane localization.

Finally, our study highlights the broader significance of lipid signaling in cellular stress responses. Identifying PS as a specific signal for HSPA1A’s localization invites exploration into how lipid changes regulate other chaperones or stress-responsive proteins. This intersection of lipidomics and proteomics offers new opportunities to investigate lipid signaling in cancer, neurodegenerative diseases, and viral infections (23,72,73).

In summary, our study identifies PS as a critical regulator of HSPA1A’s PM localization during stress, advancing our understanding of the lipid-based control of chaperone function. Beyond providing insights into cellular stress biology, this work highlights therapeutic opportunities for targeting lipid-chaperone pathways in cancer and other diseases characterized by altered lipid metabolism.

## Abbreviations

ANOVA: Analysis of Variance
cDNA: Complementary DNA
CTCF: Corrected Total Cell Fluorescence
DMSO: Dimethyl Sulfoxide
DMEM: Dulbecco’s Modified Eagle Medium
FASN: Fatty Acid Synthase
FBS: Fetal Bovine Serum
GFP: Green Fluorescent Protein
HSD: Honestly Significant Difference
HSPA1A: Heat Shock Protein A1A
LDS: Lithium Dodecyl Sulfate
MβCD: Methyl-β-Cyclodextrin
MEM: Minimum Essential Medium
MS/MS: Tandem Mass Spectrometry
NEAA: Non-Essential Amino Acids
PBS: Phosphate-Buffered Saline
PFA: Paraformaldehyde
PI4P: Phosphatidylinositol 4-Phosphate
PM: Plasma Membrane
PS: Phosphatidylserine
QTOF: Quadrupole Time-of-Flight
RFP: Red Fluorescent Protein
RNAi: RNA Interference
RIPA: Radioimmunoprecipitation Assay
RT: Room Temperature
SDS-PAGE: Sodium Dodecyl Sulfate-Polyacrylamide Gel Electrophoresis
siRNA: Small Interfering RNA;
WGA: Wheat Germ Agglutinin.

## References Cited

1. Bukau, B., and Horwich, A. L. (1998) The Hsp70 and Hsp60 chaperone machines. Cell 92, 351–366

2. Lindquist, S., and Craig, E. A. (1988) The heat-shock proteins. Annu Rev Genet 22, 631–677

3. Balogi, Z., Multhoff, G., Jensen, T. K., Lloyd-Evans, E., Yamashima, T., Jaattela, M., Harwood, J. L., and Vigh, L. (2019) Hsp70 interactions with membrane lipids regulate cellular functions in health and disease. Prog Lipid Res 74, 18–30

4. Craig, E. A. (2018) Hsp70 at the membrane: driving protein translocation. BMC Biol 16, 11

5. De Maio, A., and Hightower, L. (2021) The interaction of heat shock proteins with cellular membranes: a historical perspective. Cell Stress Chaperones 26, 769–783

6. Gehrmann, M., Liebisch, G., Schmitz, G., Anderson, R., Steinem, C., De Maio, A., Pockley, G., and Multhoff, G. (2008) Tumor-specific Hsp70 plasma membrane localization is enabled by the glycosphingolipid Gb3. PLoS ONE 3, e1925

7. Multhoff, G. (2007) Heat shock protein 70 (Hsp70): membrane location, export and immunological relevance. Methods 43, 229–237

8. Multhoff, G., and Hightower, L. E. (1996) Cell surface expression of heat shock proteins and the immune response. Cell Stress Chaperones 1, 167–176

9. Schilling, D., Gehrmann, M., Steinem, C., De Maio, A., Pockley, A. G., Abend, M., Molls, M., and Multhoff, G. (2009) Binding of heat shock protein 70 to extracellular phosphatidylserine promotes killing of normoxic and hypoxic tumor cells. Faseb J 23, 2467–2477

10. Vega, V. L., Rodriguez-Silva, M., Frey, T., Gehrmann, M., Diaz, J. C., Steinem, C., Multhoff, G., Arispe, N., and De Maio, A. (2008) Hsp70 translocates into the plasma membrane after stress and is released into the extracellular environment in a membrane-associated form that activates macrophages. J Immunol 180, 4299–4307

11. Lamprecht, C., Gehrmann, M., Madl, J., Romer, W., Multhoff, G., and Ebner, A. (2018) Molecular AFM imaging of Hsp70-1A association with dipalmitoyl phosphatidylserine reveals membrane blebbing in the presence of cholesterol. Cell Stress Chaperones 23, 673–683

12. Arispe, N., Doh, M., Simakova, O., Kurganov, B., and De Maio, A. (2004) Hsc70 and Hsp70 interact with phosphatidylserine on the surface of PC12 cells resulting in a decrease of viability. FASEB J 18, 1636–1645

13. Armijo, G., Okerblom, J., Cauvi, D. M., Lopez, V., Schlamadinger, D. E., Kim, J., Arispe, N., and De Maio, A. (2014) Interaction of heat shock protein 70 with membranes depends on the lipid environment. Cell Stress Chaperones

14. Bilog, A. D., Smulders, L., Oliverio, R., Labanieh, C., Zapanta, J., Stahelin, R. V., and Nikolaidis, N. (2019) Membrane Localization of HspA1A, a Stress Inducible 70-kDa Heat-Shock Protein, Depends on Its Interaction with Intracellular Phosphatidylserine. Biomolecules 9, 152

15. McCallister, C., Kdeiss, B., and Nikolaidis, N. (2015) HspA1A, a 70-kDa heat shock protein, differentially interacts with anionic lipids. Biochem Biophys Res Commun

16. McCallister, C., Kdeiss, B., and Nikolaidis, N. (2016) Biochemical characterization of the interaction between HspA1A and phospholipids. Cell Stress Chaperones 21, 41–53

17. McCallister, C., Kdeiss, B., Oliverio, R., and Nikolaidis, N. (2016) Characterization of the binding between a 70-kDa heat shock protein, HspA1A, and phosphoinositides. Biochem Biophys Res Commun 472, 270–275

18. McCallister, C., Siracusa, M. C., Shirazi, F., Chalkia, D., and Nikolaidis, N. (2015) Functional diversification and specialization of cytosolic 70-kDa heat shock proteins. Sci Rep 5, 9363

19. Smulders, L., Altman, R., Briseno, C., Saatchi, A., Wallace, L., AlSebaye, M., Stahelin, R. V., and Nikolaidis, N. (2022) Phosphatidylinositol Monophosphates Regulate the Membrane Localization of HSPA1A, a Stress-Inducible 70-kDa Heat Shock Protein. Biomolecules 12

20. Tagaeva, R., Efimova, S., Ischenko, A., Zhakhov, A., Shevtsov, M., and Ostroumova, O. (2023) A new look at Hsp70 activity in phosphatidylserine-enriched membranes: chaperone-induced quasi-interdigitated lipid phase. Sci Rep 13, 19233

21. Makky, A., Czajor, J., Konovalov, O., Zhakhov, A., Ischenko, A., Behl, A., Singh, S., Abuillan, W., and Shevtsov, M. (2023) X-ray reflectivity study of the heat shock protein Hsp70 interaction with an artificial cell membrane model. Sci Rep 13, 19157

22. Somu, P., Mohanty, S., Basavegowda, N., Yadav, A. K., Paul, S., and Baek, K. H. (2024) The Interplay between Heat Shock Proteins and Cancer Pathogenesis: A Novel Strategy for Cancer Therapeutics. Cancers (Basel) 16

23. Beloribi-Djefaflia, S., Vasseur, S., and Guillaumond, F. (2016) Lipid metabolic reprogramming in cancer cells. Oncogenesis 5, e189

24. Corbet, C., and Feron, O. (2017) Emerging roles of lipid metabolism in cancer progression. Curr Opin Clin Nutr Metab Care 20, 254–260

25. Vasseur, S., and Guillaumond, F. (2022) Lipids in cancer: a global view of the contribution of lipid pathways to metastatic formation and treatment resistance. Oncogenesis 11, 46

26. Balogh, G., Peter, M., Liebisch, G., Horvath, I., Torok, Z., Nagy, E., Maslyanko, A., Benko, S., Schmitz, G., Harwood, J. L., and Vigh, L. (2010) Lipidomics reveals membrane lipid remodeling and release of potential lipid mediators during early stress responses in a murine melanoma cell line. Biochim Biophys Acta 1801, 1036–1047

27. Peksel, B., Gombos, I., Peter, M., Vigh, L., Jr., Tiszlavicz, A., Brameshuber, M., Balogh, G., Schutz, G. J., Horvath, I., Vigh, L., and Torok, Z. (2017) Mild heat induces a distinct “eustress” response in Chinese Hamster Ovary cells but does not induce heat shock protein synthesis. Sci Rep 7, 15643

28. McCallister, C., Kdeiss, B., and Nikolaidis, N. (2015) Biochemical characterization of the interaction between HspA1A and phospholipids. Cell Stress Chaperones

29. Sahu, R., Kaushik, S., Clement, C. C., Cannizzo, E. S., Scharf, B., Follenzi, A., Potolicchio, I., Nieves, E., Cuervo, A. M., and Santambrogio, L. (2011) Microautophagy of cytosolic proteins by late endosomes. Developmental cell 20, 131–139

30. De Maio, A., and Hightower, L. E. (2021) Heat shock proteins and the biogenesis of cellular membranes. Cell Stress Chaperones 26, 15–18

31. Nimmervoll, B., Chtcheglova, L. A., Juhasz, K., Cremades, N., Aprile, F. A., Sonnleitner, A., Hinterdorfer, P., Vigh, L., Preiner, J., and Balogi, Z. (2015) Cell surface localised Hsp70 is a cancer specific regulator of clathrin-independent endocytosis. FEBS Lett 589, 2747–2753

32. Hess, K., Oliverio, R., Nguyen, P., Le, D., Ellis, J., Kdeiss, B., Ord, S., Chalkia, D., and Nikolaidis, N. (2018) Concurrent action of purifying selection and gene conversion results in extreme conservation of the major stress-inducible Hsp70 genes in mammals. Sci Rep 8, 5082

33. Yeung, T., Gilbert, G. E., Shi, J., Silvius, J., Kapus, A., and Grinstein, S. (2008) Membrane phosphatidylserine regulates surface charge and protein localization. Science 319, 210–213

34. Cho, K. J., van der Hoeven, D., Zhou, Y., Maekawa, M., Ma, X., Chen, W., Fairn, G. D., and Hancock, J. F. (2016) Inhibition of Acid Sphingomyelinase Depletes Cellular Phosphatidylserine and Mislocalizes K-Ras from the Plasma Membrane. Mol Cell Biol 36, 363–374

35. Husby, M. L., Amiar, S., Prugar, L. I., David, E. A., Plescia, C. B., Huie, K. E., Brannan, J. M., Dye, J. M., Pienaar, E., and Stahelin, R. V. (2022) Phosphatidylserine clustering by the Ebola virus matrix protein is a critical step in viral budding. EMBO Rep 23, e51709

36. Kim, W., Deik, A., Gonzalez, C., Gonzalez, M. E., Fu, F., Ferrari, M., Churchhouse, C. L., Florez, J. C., Jacobs, S. B. R., Clish, C. B., and Rhee, E. P. (2019) Polyunsaturated Fatty Acid Desaturation Is a Mechanism for Glycolytic NAD(+) Recycling. Cell Metab 29, 856–870 e857

37. Levin, G., Duffin, K. L., Obukowicz, M. G., Hummert, S. L., Fujiwara, H., Needleman, P., and Raz, A. (2002) Differential metabolism of dihomo-gamma-linolenic acid and arachidonic acid by cyclo-oxygenase-1 and cyclo-oxygenase-2: implications for cellular synthesis of prostaglandin E1 and prostaglandin E2. Biochem J 365, 489–496

38. Zhang, L., Ramtohul, Y., Gagne, S., Styhler, A., Wang, H., Guay, J., and Huang, Z. (2010) A multiplexed cell assay in HepG2 cells for the identification of delta-5, delta-6, and delta-9 desaturase and elongase inhibitors. J Biomol Screen 15, 169–176

39. Li, J., Condello, S., Thomes-Pepin, J., Ma, X., Xia, Y., Hurley, T. D., Matei, D., and Cheng, J. X. (2017) Lipid Desaturation Is a Metabolic Marker and Therapeutic Target of Ovarian Cancer Stem Cells. Cell Stem Cell 20, 303–314 e305

40. Chen, L. Y., Wu, D. S., and Shen, Y. A. (2024) Fatty acid synthase inhibitor cerulenin hinders liver cancer stem cell properties through FASN/APP axis as novel therapeutic strategies. J Lipid Res 65, 100660

41. Mahammad, S., and Parmryd, I. (2015) Cholesterol depletion using methyl-beta-cyclodextrin. *Methods in molecular biology (Clifton*, N.J 1232, 91–102

42. Collins, T. J. (2007) ImageJ for microscopy. Biotechniques 43, 25–30

43. Burgess, A., Vigneron, S., Brioudes, E., Labbe, J. C., Lorca, T., and Castro, A. (2010) Loss of human Greatwall results in G2 arrest and multiple mitotic defects due to deregulation of the cyclin B-Cdc2/PP2A balance. Proceedings of the National Academy of Sciences of the United States of America 107, 12564–12569

44. Johnson, K. A., Taghon, G. J., Scott, J. L., and Stahelin, R. V. (2016) The Ebola Virus matrix protein, VP40, requires phosphatidylinositol 4,5-bisphosphate (PI(4,5)P2) for extensive oligomerization at the plasma membrane and viral egress. Sci Rep 6, 19125

45. Bilog, A. D., Smulders, L., Oliverio, R., Labanieh, C., Zapanta, J., Stahelin, R. V., and Nikolaidis, N. (2019) Membrane Localization of HspA1A, a Stress Inducible 70-kDa Heat-Shock Protein, Depends on Its Interaction with Intracellular Phosphatidylserine. Biomolecules 9

46. Oliverio, R., Nguyen, P., Kdeiss, B., Ord, S., Daniels, A. J., and Nikolaidis, N. (2018) Functional characterization of natural variants found on the major stress inducible 70-kDa heat shock gene, HSPA1A, in humans. Biochem Biophys Res Commun 506, 799-804

47. Cajka, T., and Fiehn, O. (2017) LC-MS-Based Lipidomics and Automated Identification of Lipids Using the LipidBlast In-Silico MS/MS Library. *Methods in molecular biology (Clifton*, N.J 1609, 149–170

48. Cajka, T., and Fiehn, O. (2016) Toward Merging Untargeted and Targeted Methods in Mass Spectrometry-Based Metabolomics and Lipidomics. Anal Chem 88, 524–545

49. Zhang, Y., Fan, S., Wohlgemuth, G., and Fiehn, O. (2023) Denoising Autoencoder Normalization for Large-Scale Untargeted Metabolomics by Gas Chromatography-Mass Spectrometry. Metabolites 13

50. Ismail, I. T., Showalter, M. R., and Fiehn, O. (2019) Inborn Errors of Metabolism in the Era of Untargeted Metabolomics and Lipidomics. Metabolites 9

51. Bowden, J. A., Heckert, A., Ulmer, C. Z., Jones, C. M., Koelmel, J. P., Abdullah, L., Ahonen, L., Alnouti, Y., Armando, A. M., Asara, J. M., Bamba, T., Barr, J. R., Bergquist, J., Borchers, C. H., Brandsma, J., Breitkopf, S. B., Cajka, T., Cazenave-Gassiot, A., Checa, A., Cinel, M. A., Colas, R. A., Cremers, S., Dennis, E. A., Evans, J. E., Fauland, A., Fiehn, O., Gardner, M. S., Garrett, T. J., Gotlinger, K. H., Han, J., Huang, Y., Neo, A. H., Hyotylainen, T., Izumi, Y., Jiang, H., Jiang, H., Jiang, J., Kachman, M., Kiyonami, R., Klavins, K., Klose, C., Kofeler, H. C., Kolmert, J., Koal, T., Koster, G., Kuklenyik, Z., Kurland, I. J., Leadley, M., Lin, K., Maddipati, K. R., McDougall, D., Meikle, P. J., Mellett, N. A., Monnin, C., Moseley, M. A., Nandakumar, R., Oresic, M., Patterson, R., Peake, D., Pierce, J. S., Post, M., Postle, A. D., Pugh, R., Qiu, Y., Quehenberger, O., Ramrup, P., Rees, J., Rembiesa, B., Reynaud, D., Roth, M. R., Sales, S., Schuhmann, K., Schwartzman, M. L., Serhan, C. N., Shevchenko, A., Somerville, S. E., St John-Williams, L., Surma, M. A., Takeda, H., Thakare, R., Thompson, J. W., Torta, F., Triebl, A., Trotzmuller, M., Ubhayasekera, S. J. K., Vuckovic, D., Weir, J. M., Welti, R., Wenk, M. R., Wheelock, C. E., Yao, L., Yuan, M., Zhao, X. H., and Zhou, S. (2017) Harmonizing lipidomics: NIST interlaboratory comparison exercise for lipidomics using SRM 1950-Metabolites in Frozen Human Plasma. J Lipid Res 58, 2275–2288

52. Chong, J., Wishart, D. S., and Xia, J. (2019) Using MetaboAnalyst 4.0 for Comprehensive and Integrative Metabolomics Data Analysis. Curr Protoc Bioinformatics 68, e86

53. Spitzer, M., Wildenhain, J., Rappsilber, J., and Tyers, M. (2014) BoxPlotR: a web tool for generation of box plots. Nat Methods 11, 121–122

54. Balogh, G., Peter, M., Glatz, A., Gombos, I., Torok, Z., Horvath, I., Harwood, J. L., and Vigh, L. (2013) Key role of lipids in heat stress management. FEBS Lett 587, 1970–1980

55. Torok, Z., Crul, T., Maresca, B., Schutz, G. J., Viana, F., Dindia, L., Piotto, S., Brameshuber, M., Balogh, G., Peter, M., Porta, A., Trapani, A., Gombos, I., Glatz, A., Gungor, B., Peksel, B., Vigh, L., Jr., Csoboz, B., Horvath, I., Vijayan, M. M., Hooper, P. L., Harwood, J. L., and Vigh, L. (2014) Plasma membranes as heat stress sensors: from lipid-controlled molecular switches to therapeutic applications. Biochim Biophys Acta 1838, 1594–1618

56. Horvath, I., Glatz, A., Nakamoto, H., Mishkind, M. L., Munnik, T., Saidi, Y., Goloubinoff, P., Harwood, J. L., and Vigh, L. (2012) Heat shock response in photosynthetic organisms: membrane and lipid connections. Prog Lipid Res 51, 208–220

57. Vigh, L., Horvath, I., Maresca, B., and Harwood, J. L. (2007) Can the stress protein response be controlled by ‘membrane-lipid therapy’? Trends in biochemical sciences 32, 357–363

58. Birge, R. B., Boeltz, S., Kumar, S., Carlson, J., Wanderley, J., Calianese, D., Barcinski, M., Brekken, R. A., Huang, X., Hutchins, J. T., Freimark, B., Empig, C., Mercer, J., Schroit, A. J., Schett, G., and Herrmann, M. (2016) Phosphatidylserine is a global immunosuppressive signal in efferocytosis, infectious disease, and cancer. Cell Death Differ 23, 962–978

59. Segawa, K., Kurata, S., Yanagihashi, Y., Brummelkamp, T. R., Matsuda, F., and Nagata, S. (2014) Caspase-mediated cleavage of phospholipid flippase for apoptotic phosphatidylserine exposure. Science 344, 1164–1168

60. Csoboz, B., Balogh, G. E., Kusz, E., Gombos, I., Peter, M., Crul, T., Gungor, B., Haracska, L., Bogdanovics, G., Torok, Z., Horvath, I., and Vigh, L. (2013) Membrane fluidity matters: hyperthermia from the aspects of lipids and membranes. Int J Hyperthermia 29, 491–499

61. Nagy, E., Balogi, Z., Gombos, I., Akerfelt, M., Bjorkbom, A., Balogh, G., Torok, Z., Maslyanko, A., Fiszer-Kierzkowska, A., Lisowska, K., Slotte, P. J., Sistonen, L., Horvath, I., and Vigh, L. (2007) Hyperfluidization-coupled membrane microdomain reorganization is linked to activation of the heat shock response in a murine melanoma cell line. Proc Natl Acad Sci U S A 104, 7945–7950

62. Escriba, P. V., Gonzalez-Ros, J. M., Goni, F. M., Kinnunen, P. K., Vigh, L., Sanchez-Magraner, L., Fernandez, A. M., Busquets, X., Horvath, I., and Barcelo-Coblijn, G. (2008) Membranes: a meeting point for lipids, proteins and therapies. J Cell Mol Med 12, 829–875

63. Smulders, L., Daniels, A. J., Plescia, C. B., Berger, D., Stahelin, R. V., and Nikolaidis, N. (2020) Characterization of the Relationship between the Chaperone and Lipid-Binding Functions of the 70-kDa Heat-Shock Protein, HspA1A. Int J Mol Sci 21

64. Albakova, Z., Armeev, G. A., Kanevskiy, L. M., Kovalenko, E. I., and Sapozhnikov, A. M. (2020) HSP70 Multi-Functionality in Cancer. Cells 9

65. Calderwood, S. K., Khaleque, M. A., Sawyer, D. B., and Ciocca, D. R. (2006) Heat shock proteins in cancer: chaperones of tumorigenesis. Trends in biochemical sciences 31, 164–172

66. Rohrig, F., and Schulze, A. (2016) The multifaceted roles of fatty acid synthesis in cancer. Nat Rev Cancer 16, 732–749

67. Horvath, I., Multhoff, G., Sonnleitner, A., and Vigh, L. (2008) Membrane-associated stress proteins: more than simply chaperones. Biochim Biophys Acta 1778, 1653–1664

68. Sherman, M., and Multhoff, G. (2007) Heat shock proteins in cancer. Ann N Y Acad Sci 1113, 192–201

69. Hu, B., Liu, G., Zhao, K., and Zhang, G. (2024) Diversity of extracellular HSP70 in cancer: advancing from a molecular biomarker to a novel therapeutic target. Front Oncol 14, 1388999

70. Murakami, N., Kuhnel, A., Schmid, T. E., Ilicic, K., Stangl, S., Braun, I. S., Gehrmann, M., Molls, M., Itami, J., and Multhoff, G. (2015) Role of membrane Hsp70 in radiation sensitivity of tumor cells. Radiat Oncol 10, 149

71. Rysman, E., Brusselmans, K., Scheys, K., Timmermans, L., Derua, R., Munck, S., Van Veldhoven, P. P., Waltregny, D., Daniels, V. W., Machiels, J., Vanderhoydonc, F., Smans, K., Waelkens, E., Verhoeven, G., and Swinnen, J. V. (2010) De novo lipogenesis protects cancer cells from free radicals and chemotherapeutics by promoting membrane lipid saturation. Cancer Res 70, 8117–8126

72. Haughey, N. J. (2010) Sphingolipids in neurodegeneration. Neuromolecular Med 12, 301–305

73. Heaton, N. S., and Randall, G. (2011) Multifaceted roles for lipids in viral infection. Trends Microbiol 19, 368–375

