## supplemental figures for "Heat-Induced Phosphatidylserine Changes Drive HSPA1A’s Plasma Membrane Localization"

### Title

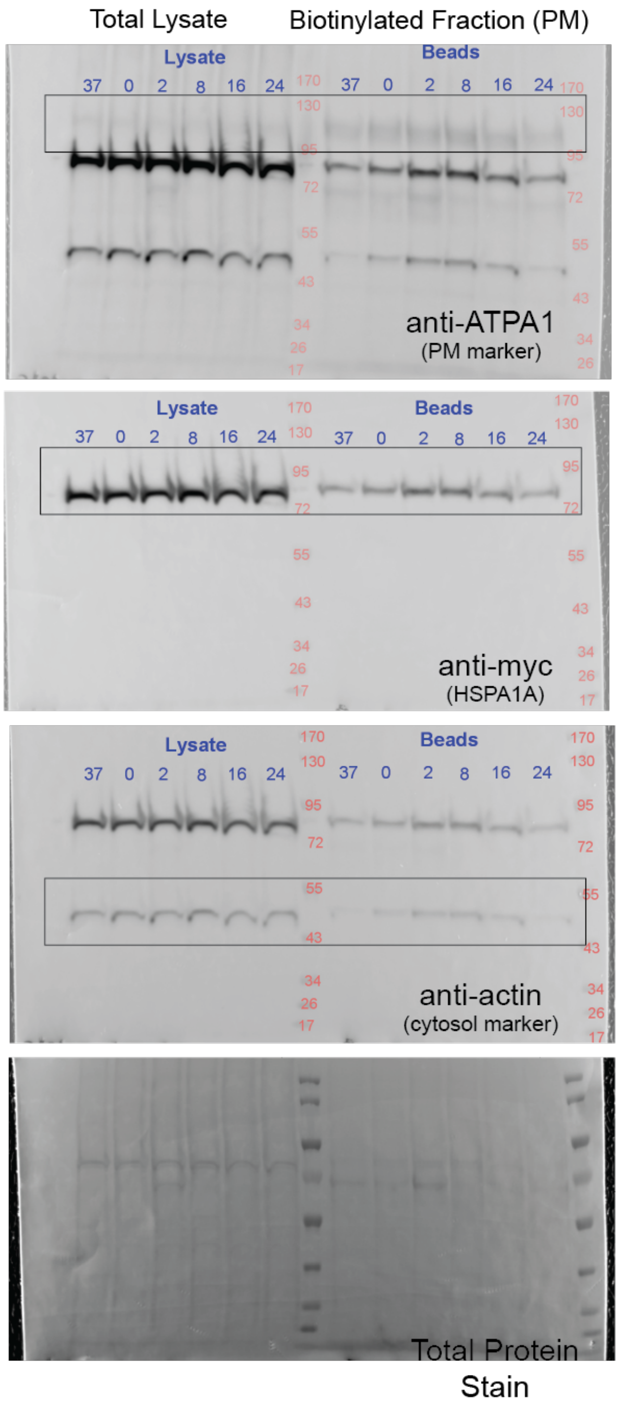


**Supplemental Figure 1. Complete Western blots showing HSPA1A's increased plasma membrane (PM) localization during recovery from mild heat shock.** The uncropped Western blots correspond to the cell surface biotinylation experiments shown in Figure 1. From top to bottom, the blots were probed with ATP1A1, myc, and actin antibodies. Molecular size markers (Fisher BioReagents™ EZ-Run™ Prestained Rec Protein Ladder) are indicated within each blot. The blots are superimposed with their corresponding white-light images to display the marker positions. The bottom panel shows total protein staining, performed using the Pierce™ Reversible Protein Stain (Thermo Scientific™) after antibody incubation.

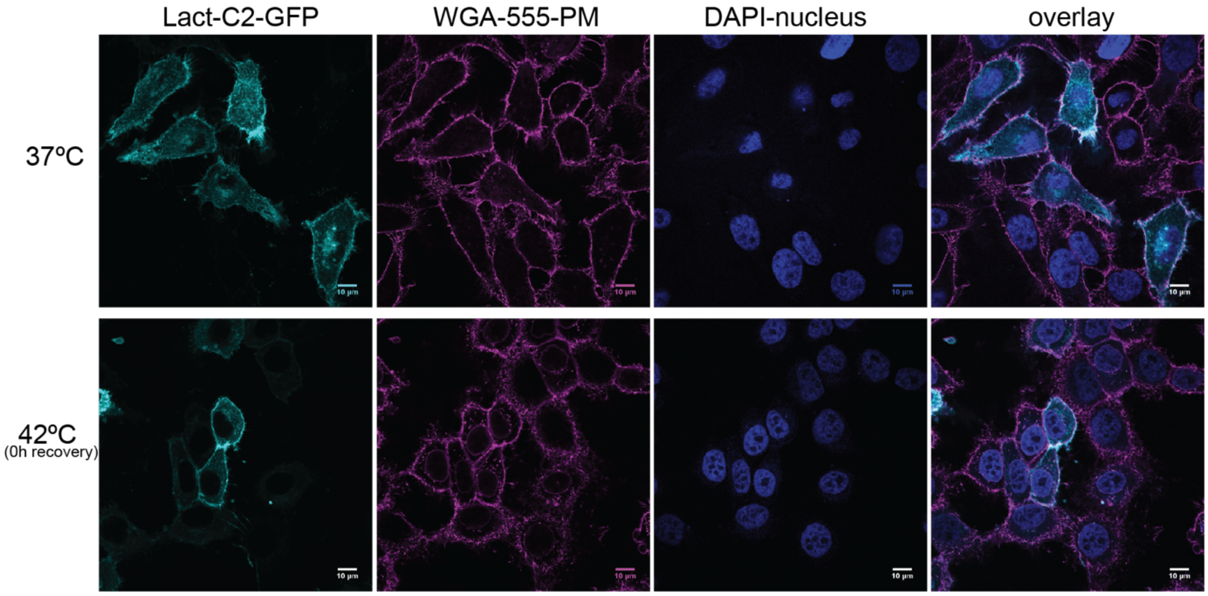


**Supplemental Figure 2. Heat shock increases phosphatidylserine (PS) levels at the plasma membrane (PM) as shown by Lact-C2 localization.** Additional confocal images of HeLa cells expressing GFP-Lact-C2, a PS biosensor. Cells were stained with WGA-FA555 to label the PM and DAPI to label nuclei. Images depict Lact-C2 localization under control conditions (37°C; top row) and immediately following heat shock (1 h at 42°C; no recovery; bottom row). Scale bar = 10 μm.

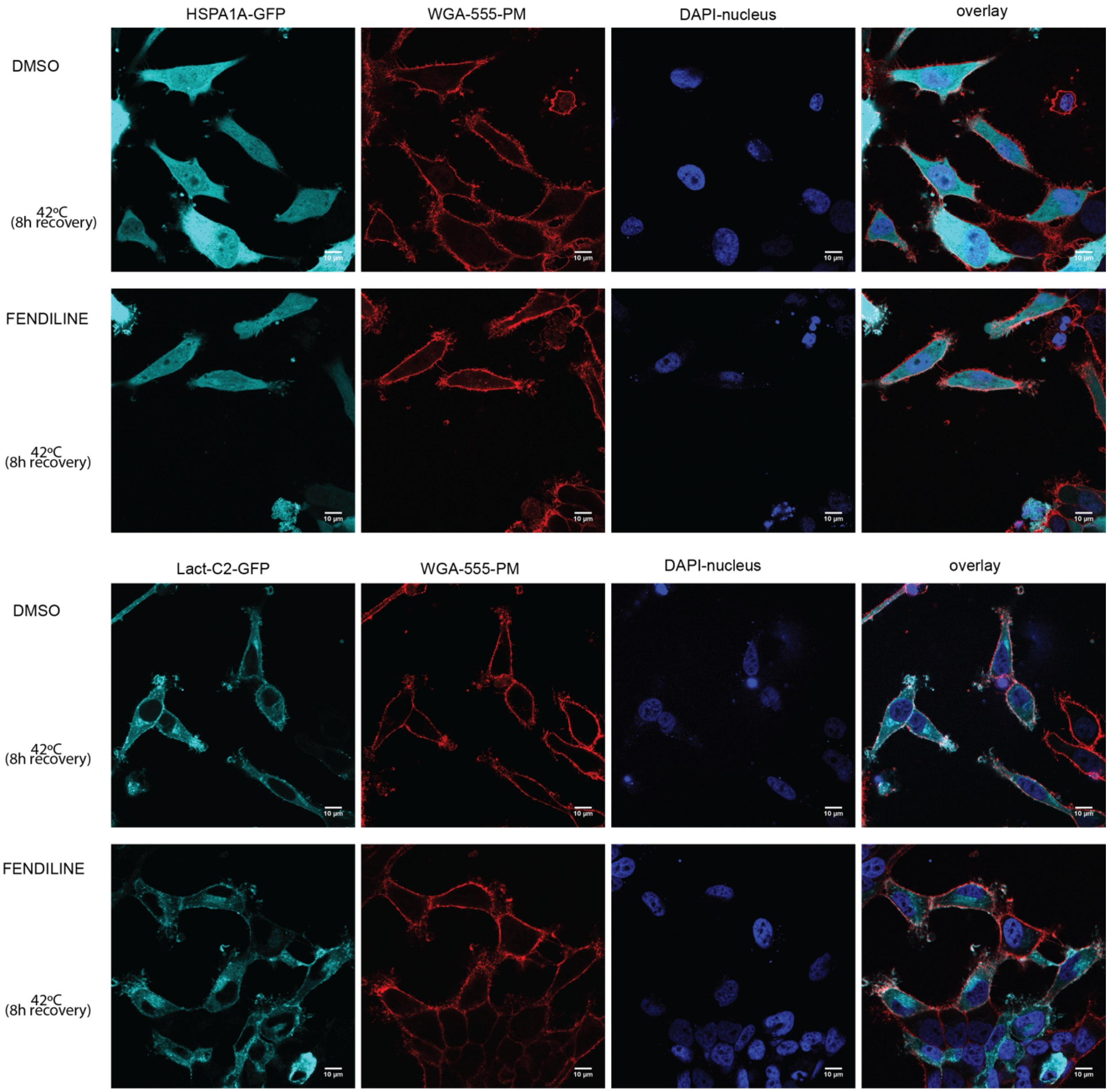


**Supplemental Figure 3. Fendiline treatment significantly decreases HSPA1A's plasma membrane (PM) localization after heat shock.** Additional confocal images of HeLa cells expressing GFP-HSPA1A (top two panels) or GFP-Lact-C2 (bottom two panels). Cells were either untreated (DMSO; top rows in each panel) or treated with 10 μM fendiline for 48 hours (bottom rows in each panel). Images depict cells after heat shock (1 h at 42°C) followed by 8 hours of recovery. The PM was stained with WGA-FA555, and nuclei were stained with DAPI (second and third images from the left in each panel). Scale bar = 10 μm.

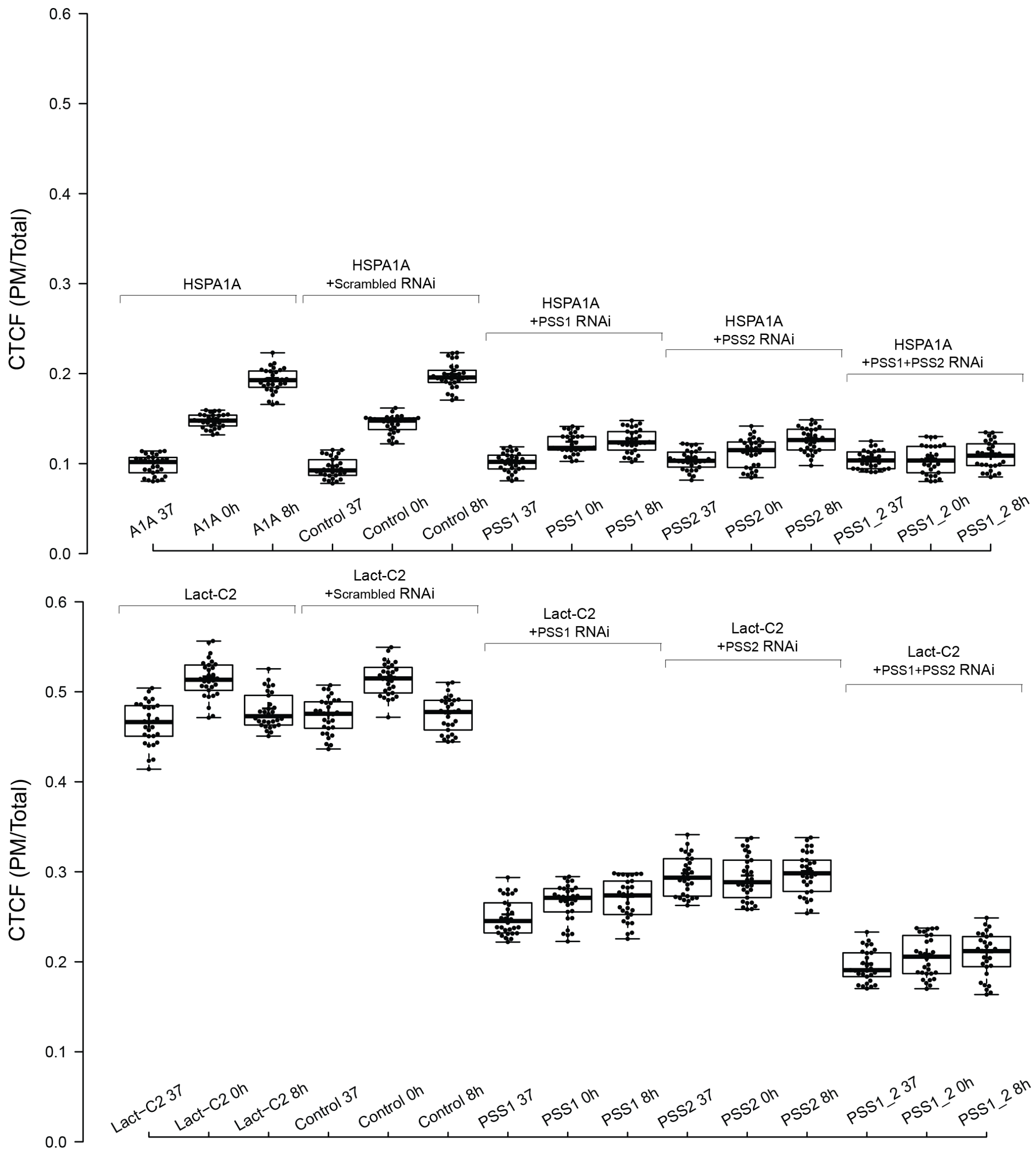


**Supplemental Figure 4. Silencing PTDSS1 and PTDSS2 genes using RNAi significantly decreases HSPA1A's plasma membrane (PM) localization.** Quantification of corrected total cell fluorescence (CTCF) as the ratio of GFP fluorescence at the PM to fluorescence in the rest of the cell. The top panel shows HSPA1A localization, while the bottom panel depicts Lact-C2 (PS biosensor) localization. Measurements were taken under control conditions (37°C) and after heat shock (1 h at 42°C), immediately following heat shock (0 hours of recovery) and after 8 hours of recovery, without RNAi, with control (scrambled RNAi), and with individual or simultaneous silencing of PTDSS1 and PTDSS2. The experiment was repeated three times, with 30 cells per condition (shown as closed circles). Box limits indicate the 25th and 75th percentiles, whiskers extend 1.5 times the interquartile range, and crosses represent sample means.
